# Intra-individual variation in leaf microbiota matches within-crown environmental heterogeneity and promotes tree performance

**DOI:** 10.64898/2026.07.24.740613

**Authors:** Lucy Saueressig, Annabell R. Wagner, Anjaharinony A.N.A. Rakotomalala, Susanne Walden, Anke Becker, Jörg Bendix, Nina Farwig, Fanhao Kong, Alexander Lach, Christian Lampei, Lars Opgenoorth, Stefan Pinkert, Tobias Trauden, Stefan Janssen, Hamed Azarbad, Robert R. Junker

## Abstract

Plant-associated microbial communities exhibit pronounced specificity across biological and spatial scales. While the patterns and accompanied functions have been well documented across and within plant species, the functional importance of intra-individual variation remains underexplored. Particularly in trees that experience strong environmental gradients within single crowns, *stratum*-specific microbiota may significantly contribute to plant performance. We experimentally tested whether variation in microbiota within the crown of *Quercus robur* is related to host performance. In mesocosm experiments, we transferred microbial communities derived from sun and shade leaves to germ-reduced clonal individuals of the same species and applied UV radiation simulating conditions that matched or mismatched the origin of the microbial inoculum (environmental matching). Our results demonstrate that matching microbiota-environment combinations increased plant performance compared to mismatching combinations. We infer that pronounced intra-individual variation of leaf-associated microbial communities not only reflects environmental heterogeneity along canopy *strata* but is functionally relevant for the plant host.

## Introduction

Phyllosphere microbiota exhibit strong heterogeneity on different organizational scales, ranging from landscapes to plant-individuals (Vorholt, 2012; Vacher *et al*., 2016; Remus-Emsermann and Schlechter, 2018). This variation reflects host-genetic as well as environmental drivers, whose relative importance may vary across scales (Laforest-Lapointe, Messier and Kembel, 2016b). On an intra-individual scale, different plant organs and tissues have been shown to be associated with distinct microbial communities (Ottesen *et al*., 2013; Junker and Keller, 2015; Cregger *et al*., 2018; Arnold *et al*., 2025; Zhang *et al*., 2025). On top of that, intra-individual variability even occurs within plant organs (e.g. leaves) and is particularly prominent in trees that exhibit a strong environmental heterogeneity within individual tree crowns (Laforest-Lapointe, Messier and Kembel, 2016b; Stone and Jackson, 2019; Herrmann *et al*., 2021).

Epiphytic leaf-associated microbial communities strongly contribute to the host’s functionality and health (Lindow and Brandl, 2003; Vorholt, 2012; Bulgarelli *et al*., 2013; Compant *et al*., 2019). For instance, phyllospheric epiphytes influence key ecological processes such as nutrient cycling, suppression of foliar pathogens, and tolerance to biotic and abiotic stressors, thereby increasing plant performance (Rastogi, Coaker and Leveau, 2013; Berg *et al*., 2014; Vacher *et al*., 2016; Villano *et al*., 2025), which has also been shown for trees (Laforest-Lapointe *et al*., 2017; Gao *et al*., 2025). Importantly, these microbiome-mediated effects likely depend not only on the mere presence of microbial strains but on their community composition, abundance and functional potential (Vacher *et al*., 2016; Lajoie, Maglione and Kembel, 2020; Sohrabi *et al*., 2023), suggesting that variation in leaf microbiota may translate into functional differences between leaves on different positions within the crown. However, it remains unknown whether intra-individual variation in leaf-associated microbiota in trees has functional consequences or whether it is merely reflecting the variability in (a)biotic conditions at different plant parts along canopy *strata*.

Within the canopy of individual trees abiotic factors vary significantly due to pronounced vertical stratification (Murakami *et al*., 2022). The upper crown is characterized by higher temperatures, light intensities and low humidity, whereas the lower tree canopy exhibits colder, more humid and shaded conditions (De Frenne *et al*., 2019; Shaik, Jallu and Doctor, 2023; Vinod *et al*., 2023). As a result, sun and shade leaves strongly differ in their morphology and physiology providing variable habitats for associated organisms (Givnish, 1988; Murakami *et al*., 2022; Wang *et al*., 2024). Herrera (2017) emphasized that such intra-individual variation in functional traits of individual organs, e.g. leaves, is ecologically relevant and represents an important level at which organismal interactions are structured within plants. Extending this concept to the phyllosphere microbiota, microbial community composition on leaves similarly varies within individual tree crowns in response to trait- and environment-driven niche differentiation (Laforest-Lapointe, Messier and Kembel, 2016b; Morella *et al*., 2020; Herrmann *et al*., 2021). Thereby, abiotic conditions may shape plant-associated microbiota both directly by influencing microbial colonization and establishment and indirectly mediated via leaf functional traits that define niche availability (Sohrabi *et al*., 2023; O’Rourke *et al*., 2025).

One very prominent abiotic factor that differs within the tree canopy is UV radiation (Brown, Parker and Posner, 1994; De Frenne *et al*., 2021). Although high doses of UV radiation can damage DNA, proteins and promote the accumulation of reactive oxygen species (ROS) in both plants and microorganisms, ambient UV radiation functions as an important environmental signal in plant-microbe interactions (Ballaré *et al*., 2011; Carvalho and Castillo, 2018; Vanhaelewyn *et al*., 2020). In plants, UV-B perception through the UVR8 receptor modifies leaf chemistry, metabolite composition and photoprotective traits, whereas microorganisms respond to UV exposure by activating DNA repair pathways, producing extracellular polymeric substance (EPS), and also photoprotective pigments (Jenkins, 2014, 2017; Carvalho and Castillo, 2018; Vanhaelewyn *et al*., 2020). Together, these direct effects on microorganisms and indirect, host-mediated changes in leaf traits influence microbial colonization and community assembly, thereby selecting for a UV-adapted microbiota (Jacobs and Sundin, 2001; Jacobs, Carroll and Sundin, 2005) that may improve host performance under subsequent UV exposure (Villano *et al*., 2025). Previous studies have demonstrated that the environment structures plant-associated microbial communities, which in turn can enhance tolerance to dynamic abiotic conditions and support plant performance (Allsup, George and Lankau, 2023; Zieschank *et al*., 2025). Accordingly, we hypothesize that variation in microbial community composition and diversity within the crowns of tree individuals on the one hand reflects variable abiotic conditions, and on the other hand also represents a mechanism that allows the trees to optimally perform despite variable environmental conditions experienced on different canopy *strata*. Thus, plant performance may be optimized by matching combinations of microbiota and environmental conditions – also on the intra-individual level. While this hypothesis has been tested on the species level (Lau and Lennon, 2012; Carrell *et al*., 2022; Allsup, George and Lankau, 2023; Zieschank *et al*., 2025), the intra-individual scale has not yet been explicitly considered in this context.

We investigated whether variation in leaf microbiota within a single tree crown is functionally relevant for the plant host. By specifically focusing on naturally structured microbiota rather than experimentally manipulated communities, this study extends existing conceptual frameworks toward a more integrated understanding of plant-microbe interactions in natural systems. As multiple biotic and abiotic factors would be confounded with intra-individual variation under natural conditions, we used a mesocosm approach with clonal individuals to test the effects of individual experimentally-controlled factors (He *et al*., 2024; Zieschank *et al*., 2025). We conducted an inoculation experiment in which leaf-associated microbiota of *Quercus robur* trees were harvested from two naturally distinct conditions in the forest (sun and shade exposed leaves) and transferred onto germ-reduced clone individuals of the same species in the laboratory (Herrmann, Munch and Buscot, 1998) (Fig. 1). Unlike seedlings, clonal individuals (from here on ramets) share an identical genetic background, allowing to attribute observed differences solely to experimental treatments. We used UV as an abiotic factor to simulate conditions that either matched or mismatched the environment from which the microbial inoculum was derived (environmental matching) (Supplementary Table 1). We hypothesized that i) the diversity and composition of leaf microbiota collected in the field differ between sun and shade leaves within individual trees, that ii) experimental conditions simulating sun and shade have comparable effects on leaf microbiota; and that iii) a match between microbiota and environmental condition positively contributes to host performance. Our results strongly support the hypothesis that environment-specific microbiota associated to different leaves within a tree individual positively affect plant performance, which suggests that intra-individual variability in microbiome composition and diversity is of ecological relevance and not simply reflects variation in the micro-environment.

**Fig. 1.**
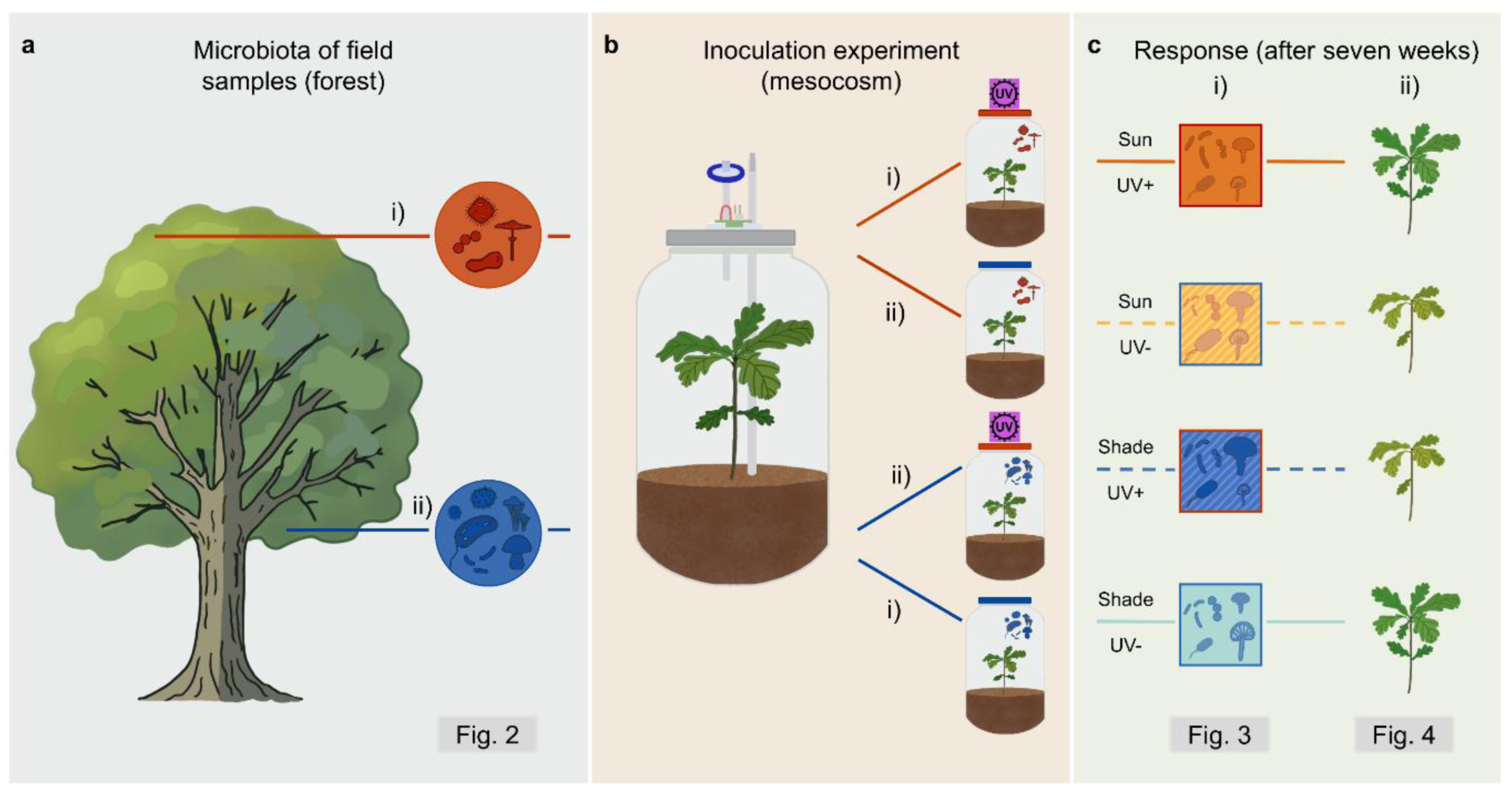
Design of the mesocosm experiment. We tested whether *stratum*-specific microbiota have a functional relevance within *Quercus robur* trees. **a**, Leaf-associated microbiota was sampled from i) sun (orange) and ii) shade (blue) positions within the canopy of *Q. robur* trees in the forest. **b**, Within a controlled mesocosm experiment, the isolated microbiota were used as microbial inocula on germ-reduced ramets of the same species. Applying a UV+ or UV-treatment, we simulated i) matching and ii) mismatching conditions according to the environment of origin of the microbial inoculum (match: sun, UV+ /shade, UV-; mismatch: sun, UV- / shade, UV+). **c**, After seven weeks, we performed high throughput 16S rRNA gene and ITS amplicon sequencing on the microbiota of treated plants and digitally phenotyped the plants to follow both i) changes in the microbial diversity and composition and ii) plant response to the applied experimental treatments. Icons were created by the authors or acquired and adapted from Phylopic.org (artists: M. Keesey, G. Dera, L. Simons).

## Material & methods

### Sampling site and leaf collection

The sampling area is located in the Marburg Open Forest (www.uni-marburg.de/mof) in the southeast of Caldern (50.84079° N, 8.68957° E). Within the sampling site, seven *Quercus robur* trees were selected and sampled from the 17^th^ to the 20^th^ of July 2023. To ensure the highest environmental contrast possible and cover within tree habitat heterogeneity, leaves were sampled at two heights integrating a vertical and horizontal gradient. Sun leaves were collected at the canopy top from south oriented branches; shade leaves were collected at the lower canopy from north oriented branches (Fig. 1a). For each canopy position three branches two leaves each were sampled resulting in *n* = 6 leaves per canopy position per tree and a total sum of *n* = 84 field samples. Leaves were collected by professional climbers using sterilized forceps and gloves to directly transfer each leaf into a petri dish within the tree. Samples were stored at 4°C until further processing in the laboratory within 3 hours after sampling. Temperature in °C and light intensity in µmol/(m²s) were captured from May to September 2023 via permanently installed dataloggers (Onset HOBO Pendant Temperature/Light datalogger UA-002-08; HOBO dataloggers, Bourne, MA, USA) at both canopy positions in each tree.

### Extracting microbiota from field samples

To isolate microbial inocula (see inoculation experiment below) of phyllospheric microorganisms, the microbiota of sun and shade-exposed leaves were collected in 20 mL PBS via high frequency vibration using a Pulsifier II (Microgen Bioproducts PUL 200 Pulsifier II™; Novacyt, Manchester, UK) for three times 30 seconds each. This method detaches microorganisms from the leaf surface while keeping damages of the plant tissue as low as possible. The microbial suspension was centrifuged in 50 mL falcon tubes for 20 minutes at 7000 RCF at 4°C. The resulting pellet was resuspended in 2 mL of the supernatant. Glycerol stocks (1:1 suspension and glycerol) were prepared and stored at – 80 °C. Prior to inoculation, the microbial suspensions (glycerol stocks) from each of the three branches were pooled per canopy position for each tree (shade microbiota *n* = 7, sun microbiota *n* = 7, total number of microbial inocula *n* = 14).

### Inoculation experiment

To assess the functional consequences of intra-individual variation in leaf microbiota, we conducted an inoculation experiment comparing matched and mismatched microbiota-environment combinations. Therefore, we combined microbiota harvested from the field samples (either from sun or shade exposed leaves) and a UV treatment mimicking the UV exposure of an average summer day in Hesse (Beckmann *et al*., 2014). Accordingly, the experiment comprised four different treatments (Fig. 1b, Supplementary Table 1). These included two matching treatments, in which ramets of the *Q. robur* clone DF159 were inoculated with sun or shade microbiota and treated with UV+ or UV-, respectively. Additionally, two mismatching treatments were included, in which ramets inoculated with sun or shade microbiota were treated with UV- and UV+, respectively (Fig. 1b, Supplementary Table 1) as well as two controls. To run the experiment under sterile, highly controlled conditions, we used a mesocosm approach. All components of the mesocosms (glass jar, lid, tubes, sealing caps, substrate, medium) were sterilized prior to usage and only handled in a sterile environment afterwards. A mesocosm consisted of a 4250 mL glass jar (Gläser und Flaschen GmbH, Wustermerk, Germany) closed with the appropriate lid. A silicon tube was inserted through a hole in the lid to water the plants without opening the jar (4 mm × 7 mm tube; Bürkle GmbH, Bellingen, Germany). The end of the tube was inserted into the substrate to avoid washing off microbes from the leaf surface while watering. The opening was closed with a 4 mm sealing cap (Reichelt electronic, Sande, Germany) throughout the experiment. Further a second, shorter 4 mm × 7 mm silicon tube attached to a syringe filter (SFCA, 25 mm diam., 0.2 µm; TH Geyer, Renningen, Germany) allowed for passive gas exchange. A schematic figure of the mesocosm setup can be found in Supplementary Fig. 1. To simulate natural UV exposure occurring in field, the lid of the jar was equipped with a customized, adjustable UV-B module UV-B LED emitting at 310 nm (narrowband ranging from 310 – 315 nm with a broad viewing angle of 120° and a flat quartz glass window ensuring Lambertian emission; model GD35R-310Z-F, 40mA, 3 mW, Roithner Lasertechnik GmbH, Vienna, Austria, for details see Supplementary Fig. 1). Using three UV-B LED chips, the UV intensity could be adjusted from 0.007 to 0.021 mW/cm². For plants which were exposed to UV, we used an irradiation intensity of 0.015 mW/cm² for 6 h and 35 minutes which equals 356 mJ/cm² per day (3.56 kJ/m²/d), matching the mean UV dosage of a sunny day in July in Hesse, Germany (Beckmann *et al*., 2014).We applied a constant intensity of 0.015 mW/cm² to reach the average daily dosage. To obtain the average daily UV-B dosage for July in Hesse, we clipped the global UV-B raster for July (Beckmann *et al*., 2014) to the Hesse administrative boundary using GIS-based spatial masking. Plants which were not exposed to UV received no UV radiation, which did not affect temperature conditions in the mesocosms. As plant material we used ramets of the *Q. robur* clone DF159 propagated in vitro under sterile conditions (Herrmann, Munch and Buscot, 1998). The oak clone therefore only carries a reduced microbiota making it a suitable organism to test effects of microbial inoculation. The ramets were planted into the mesocosms under sterile conditions at a rooted stage, using 700 g of pre-washed and sterilized clay as the substrate (Calcined Clay Drying Agent; Diamond Pro, Arlington, TX, USA). The substrate was mixed with 365 mL of ½ MS medium containing vitamins and trace elements to reach 55% of the substrate’s water holding capacity. Plants were acclimatized for three weeks prior to the start of the experiment. For microbial inoculation the stock solutions received from field samples (see above) were used as microbial inocula on the germ-reduced ramets. For inoculation, 700 µL of the microbial inoculum (pooled stock suspensions) was sprayed onto a plant using the BLUESTAR® Ecospray micro-diffusor (diam. 0.50 mm, 0.27 mL/s; Bluestar Forensic, Monaco). For controls, 1:1 PBS-glycerol mixture was used as inoculum. The substrate was covered by a layer of plastic foil attached to filter paper prior to spraying to minimize the inoculation of the soil by absorbing dripping suspension. Using a micro-diffusor for inoculation enabled homogeneous inoculation of the whole plant, including both the upper and lower side of the leaves (Fukami *et al*., 2016; Preininger *et al*., 2018). To allow sufficient time for microbial colonization of the new habitat, UV exposure was initiated one week after inoculation (Innerebner, Knief and Vorholt, 2011; Remus-Emsermann and Schlechter, 2018). Plants were grown in a climate chamber at 21.5°C with a 16 h/8 h light-dark cycle (LED plant lamp Floris N1000WN, Photosynthetic Photon Flux Density (PPFD) = 110 µmol/m²s; Neusius Pflanzenlampen, Saarbrücken, Germany) for the initial three weeks of acclimatization, one week of microbial establishment, and the experimental period of seven weeks to assess changes in the plant phenotype and the microbial response to the applied treatments (Fig. 1c). Plants were watered once with 50 mL of ½ MS medium containing vitamins and trace elements five weeks after planting. Throughout the experiment, humidity within the mesocosms was constantly at 100%. At the end of the experiment, two leaves of each ramet were used to collect microbiota from the treated plants as described above for the field samples (pulsifier approach). For comparability we used leaves of approximately the same size and position (top layer or one layer below), thereby the youngest, freshly emerged leaves were excluded.

### Digital plant phenotyping

Prior to planting and at the end of the experiment after removing the plants from mesocosms, whole plants were phenotyped using an automated plant scanner (PlantEye F500, Phenospex, Heerlen, The Netherlands) (Zieschank and Junker, 2023; Zieschank *et al*., 2025). The customized 3D plant scanner is able to capture 14 physiological and spectral plant parameters simultaneously via laser scans, making it an efficient and non-invasive tool to derive plant trait information. The device can further operate under sterile condition as it was modified to be setup under a safety bench, avoiding contaminations with microorganism once the plants were removed from the mesocosms for scanning. With sterilized forceps each plant was positioned centrally to the device to be scanned. Later, only the above ground parts of the plant were considered for analysis by setting a certain cutoff height. Scans were processed using the software HortControl (Phenospex, Heerlen, The Netherlands). In the following analysis we focused on the Normalized Difference Vegetation Index (NDVI), which is often used to infer vegetation health, biomass, and productivity (Pettorelli *et al*., 2005; Kaur *et al*., 2015) and the difference in maximum height (ΔH_max_) of a plant before and after the experiment (Pérez-Harguindeguy *et al*., 2013), which were set as proxies for plant performance. NDVI is calculated as (NIR - RED) / (NIR + RED), with NIR and RED being reflectance values in the near-infrared and red spectral bands, respectively. Values above 0.5 can be indicative of a healthy vegetation (Ipanaqué *et al*., 2025). For network analysis we further included the plant traits Greenness, Hue, Normalized Pigment Chlorophyll Ratio Index (NPCI), Plant Senescence Reflectance Index (PSRI), Difference in Digital biomass, Leaf area (LA), Leaf area index (LAI), and Light penetration depth (LPD) (Zieschank and Junker, 2023).

### DNA extraction, Amplicon Sequencing and data preparation

Total genomic DNA was extracted from sun and shade microbial inocula, obtained from field samples, and from treated plants after the inoculation experiment using the DNeasy Plant Pro Kit (Qiagen, Hilden) following the manufacturer’s protocol (adjustment: 1 mL of microbial extract used as initial input) to analyze bacterial and fungal community composition. The isolated DNA was sent to LGC Genomics (Berlin, Germany) to carry out high throughput 16S and ITS amplicon sequencing. The V5-V6 region of the bacterial 16S rRNA gene was amplified using the primer pair 799F (AACMGGATTAGATACCCKG) and 1115R (AGGGTTGCGCTCGTTG) (Thijs *et al*., 2017), whereas the fungal ITS1 region was amplified using the primer pair ITS1F (CTTGGTCATTTAGAGGAAGTAA) and ITS2 (GCTGCGTTCTTCATCGATGC) (White *et al*., 1990; Gardes and Bruns, 1993). Amplicons were sequenced with Illumina MiSeq v3 (2 × 300 bp paired-end reads) to characterize bacterial and fungal communities. The bacterial primer set was specifically designed to minimize the amplification of host plant DNA and enrich bacterial sequences (Thijs *et al*., 2017). Bioinformatic analysis were performed with QIIME 2 version 2024.10 (Bolyen *et al*., 2019). Basecalling and demultiplexing was performed by LGC Genomics via bcl2fastq version v2.20 with subsequent adapter and primer trimming via Cutadapt v5.1 (Martin, 2011). To balance read errors and taxonomic resolutions, we chose to call Amplicon Sequence Variants (ASVs) via DADA2 (q2-dada2 plugin) from the first 150 bp of forward reads only. Taxonomy of the bacteria was assigned to ASVs using the naive Bayes classifier (q2-feature-classifier plugin) against the Greengenes2 reference sequences version 2024.09 (McDonald *et al*., 2024). Taxonomy of the fungi was assigned to ASVs against the UNITE 97% reference sequences (Kõljalg *et al*., 2020). Archaeal and other eukaryotic sequences besides fungi were excluded from further analysis by subsetting the Domain or Kingdom to bacteria and fungi, respectively. Unassigned phyla were removed from the datasets. Non-target sequences of bacteria were filtered by excluding Cyanobacter and Chloroflexota at phylum level, as well as Rickettsiales and Diplorickettsiales at order level. Non-target sequences of fungi were filtered by excluding the genus Malassezia.

### Statistical analyses

All statistical analyses were performed using R version 4.5.3 (R Core Team, 2026). To ensure an appropriate quality of our sequencing results, we computed species-accumulation curves using the R package *vegan* v2.7–3 (Oksanen *et al*., 2026) and determined the number of reads required to obtain 90% of the ASV richness in each sample. The median of all samples was set as a cutoff threshold (bacteria median = 1876; fungi median = 551) for downstream analyses. For bacteria, one sample had to be removed after setting the cutoff threshold resulting in *n* = 77 samples, whereas for fungi four samples were removed resulting in *n* = 73 samples. Lastly, we filtered low frequency taxa by removing ASVs that occurred in less than two samples. To analyze the microbial alpha diversity, we accounted for differences in sequencing depth by rarefying the data to the minimum number of reads across samples (iterations *n* = 9999) before calculating alpha diversity metrics using the R package *rtk* v0.2.6.1 (Saary *et al*., 2017). For both bacterial and fungal datasets, richness and Shannon diversity indices (*H*^′^ = − ∑^*s*^_i=1_ p_i_ ln(p_i_)) were calculated separately for i) the sun and shade microbiota of the field samples (used as inocula) to examine difference according to the position within the tree canopy and ii) the microbiota of treated ramets at the end of the experiment to evaluate difference resulting from experimental treatments. Differences in diversity among groups were assessed using analysis of variance (ANOVA) followed by a post hoc Tukey’s HSD test using the R package *agricolae* v1.3-7 (Mendiburu, 2023). Differential abundance of ASVs among treatments was analyzed separately for bacteria and fungi using the R package *DESeq2* v1.40.2 (Love, Huber and Anders, 2014). ASV counts from the non-rarified datasets were modeled using negative binominal GLMs with treatment as fixed effect. Size factors were estimated using the “poscounts” method, which is suitable for sparse microbial count data, while all other *DESeq2* parameters were kept at their default values. The treatment factor comprised the two microbial inocula derived from field samples and the six treatments of the inoculation experiment with two matching and two mismatching conditions (Supplementary Table 1). Five predefined contrasts were tested pairwise to compare sun and shade microbiota of the field samples used as microbial inocula (Sun_start vs. Shade_start) and microbiota of the oak clones after the experimental treatment according to UV condition (Sun_UV+ vs. Sun_UV-; Shade_UV+ vs. Shade_UV-; Control_UV+ vs. Control_UV-). Statistical significance of differential ASV abundance between treatments was assessed using Wald tests with Benjamini-Hochberg false discovery rate (FDR) correction. Log_2_ fold changes for each contrast were visualized using volcano plots. To display overall community composition of bacteria and fungi, shared and unique ASVs across experimental groups were identified from presence-absence matrices and visualized in Upset plots and Venn diagrams. To further analyze community composition, the non-rarified ASV count data were normalized via cumulative sum scaling (CSS) as implemented in the R package *metagenomeSeq* v1.52.0 (Paulson *et al*., 2013) followed by min-max normalization. Bray-Curtis dissimilarities were calculated from the normalized data and visualized by non-metric multidimensional scaling (NMDS). Differences in community composition among treatments were tested using permutational analysis of variance (PERMANOVA; 9999 permutations) implemented in the R package *vegan* v2.7–3 (Oksanen *et al*., 2026). Taxonomic composition was visualized at the family level as bar plots of the 12 most abundant bacterial and fungal families based on mean relative abundance across experimental groups. Unclassified and remaining low-abundance families were pooled in “Unclassified” and “Others”, respectively. In order to assess abiotic variation within the canopies of the sampled forest trees, differences in temperature and light intensity were tested separately for each month using Wilcoxon rank-sum test (for temperature day and night were analyzed separately).

To estimate the plant responses to microbial inoculation and the subsequent experimental treatment (matching or mismatching conditions), we focused on two plant performance proxies, the NDVI and the difference in maximum height (ΔH_max_), reflecting growth. Therefore, we conducted a two-factorial analysis of variance (ANOVA) testing the interactive effects of inoculation type (sun or shade derived microbial inoculum) and UV exposure on plant traits (either NDVI or ΔH_max_). Post-hoc pairwise comparisons were performed using Tukey’s Honest Significance Difference (HSD) test.

We used a correlation-based network to further analyze interactions among bacteria, fungi, and the host plant phenotype. Prior to network construction, bacterial and fungal non-rarified ASV count data were normalized via cumulative sum scaling (CSS) as described above. Normalized microbial matrices were merged and aligned with plant trait data (normalized by min–max scaling). After removing controls, the dataset was partitioned into matching and mismatching conditions to enable treatment specific analyses. For each subset, pairwise associations were calculated using Spearman’s rank correlations between i) microbial taxa, ii) plant traits, and iii) microbes and plant traits. The correlations required a minimum of nine co-occurrences and excluded pairs in which one or both values were zero. Based on significant correlations (p < 0.05), networks were built and visualized using the R package *igraph* (Csárdi *et al*., 2023). To characterize the structural properties of the network, standard network metrics were calculated: number of nodes and edges (network size), network density (proportions of realized links), mean node degree (average links per node), mean node strength (average weighted connectivity based on absolute correlation values), modularity based on the Louvain clustering algorithm (strength of subdivision into modules), number of connected components (network fragmentation), and size of the largest component. Node sizes within the network visualizations were scaled according to rescaled betweenness centrality values highlighting potential keystone taxa.

## Results

### Canopy *stratum*-specific leaf microbiota within tree individuals in the forest

To characterize the associated microbiota of sun and shade leaves within the crown of *Q. robur* that were also used as inoculates for mesocosm experiments, we performed high-throughput amplicon sequencing of the 16S rRNA gene of bacteria and the fungal ITS1 region. Among shade leaves, we obtained 403 bacterial and 281 fungal ASVs (*n* = 7 trees), whereas sun leaves harbored 212 bacterial and 196 fungal ASVs (*n* =7 trees). Across all 14 samples, a total of 481 bacterial and 337 fungal ASVs were identified (Supplementary Fig. 2). Shannon diversity and richness of shade microbiota were significantly higher compared to sun microbiota for both bacteria and fungi (Shannon diversity: bacteria: ANOVA: *F*_1,12_ = 7.834, *P* = 0.016; Fig. 2a; fungi: ANOVA: *F*_1,12_ = 12.81, *P* = 0.004; Fig. 2c, Supplementary Table 2; richness: see Supplementary Fig. 3a, c, Supplementary Table 2). The composition of microbial communities differed with canopy position for both bacteria and fungi (Supplementary Fig. 4, 5). Further, the differential abundance analysis revealed that canopy position significantly drives differences in the abundance of bacterial and fungal ASVs, respectively (Fig. 2b, d). In shaded conditions 9.7% and 4.3% of the bacterial and fungal ASVs were more abundant, whereas 2.5% of bacterial and 2.3% of fungal ASVs were more abundant in sun conditions (Fig. 2b, d). Detailed taxonomic information are provided in the Supplementary Table 3a, b. The analysis of environmental data from loggers installed at same locations as the leaves from which the microbiota was retrieved, showed that microbial communities in the upper canopy were exposed to higher light intensity and temperature than communities in the lower canopy (Supplementary Fig. 6).

**Fig. 2.**
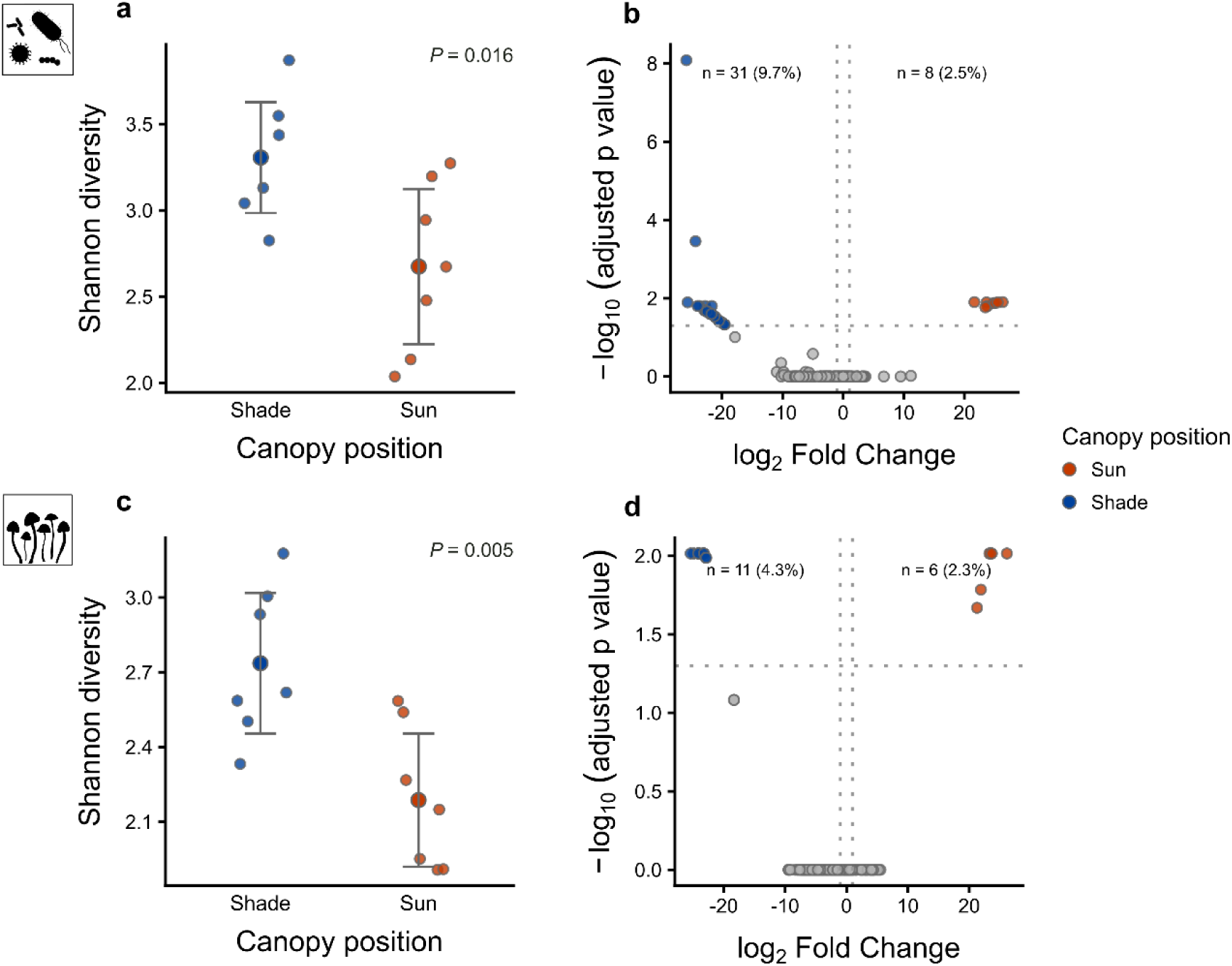
Alpha diversity and differential abundance of bacterial and fungal communities along canopy *strata*. **a**, **c**, Shannon diversity index (*H’*) of bacterial (**a**) and fungal (**c**) leaf-associated communities along canopy positions (sun, orange; shade, blue): Microbiota was samples from *n* = 7 *Q. robur* forest trees and analyzed via 16S rRNA gene (**a**) and ITS (**b**) amplicon sequencing. Large central circles and error bars indicate mean and 95% confidence intervals (CI), respectively. **b**, **d**, Volcano plots illustrating differential abundance of individual bacteria (**b**) and fungal (**d**) amplicon sequence variants (ASVs) between sun and shade canopy positions, calculated using *DESeq2*. Points represent individual ASVs plotted by their log_2_ fold change and – log_10_-transformed adjusted *P* values (*P_adj_*). Extreme log_2_ fold change values (±20) indicate ASVs with zero or near-zero values in one treatment, resulting in numerical capping by *DESeq2* (Love, Huber and Anders, 2014). Dotted lines indicate the significance thresholds (*P_adj_* < 0.05 and log_2_ fold change | > 1). Enriched ASVs are color-coded by canopy position (sun, orange; shade, blue). Icons were created by the authors or acquired and adapted from Phylopic.org (artists: F. Vaux, L. Simons). Results of the analyses of variance (ANOVA) are provided in the main text and Supplementary Table 2. For detailed taxonomic information of the significant bacterial and fungal ASVs see Supplementary Table 3a and b, respectively.

### Matching microbiota and environment promote plant performance within an inoculation experiment

To test whether *stratum*-specific microbial communities have a functional relevance, we inoculated germ-reduced oak ramets in mesocosms with microbiota derived from sun or shade leaves of forest trees as described above. To simulate sun and shade leaf conditions, oak ramets were experimentally either exposed to UV radiation or not. UV radiation in mesocosms matched intensities as observed on sun-exposed leaves in the field in summer (Beckmann *et al*., 2014). Therefore, we were able to treat oak ramets with microbiota-environment combinations that matched, i.e. sun microbiota with UV-treatment or shade microbiota without UV-treatment. As contrast, we also created mismatching combinations, i.e. sun microbiota without UV-treatment or shade microbiota with UV-treatment (Supplementary Table 1). After seven weeks, we assessed the alpha diversity and differential abundance of bacterial and fungal community members as well as the plant performance measured as Normalized Difference Vegetation Index (NDVI) and absolute growth in the experimental period (ΔH_max_).

Among treatments, 815 bacterial and 89 fungal ASVs were identified respectively (*n* = 55, *n* = 54). Within controls, 280 bacterial and 65 fungal ASVs were found (*n* = 8, *n* = 5) (Supplementary Fig. 2). Experimental treatments altered microbial community composition as compared to the field samples described above (Supplementary Fig. 4, 5) and also reduced Shannon diversity and richness (Supplementary Fig. 7, Supplementary Table 2). Within laboratory treatments, Shannon diversity and richness of both, bacteria and fungi, did not differ for any of the applied treatments (Shannon diversity: bacteria: ANOVA: *F*_3,51_ = 0.182, *P* = 0.908; Fig. 3a; fungi: ANOVA: *F*_3,50_ = 0.556, *P* = 0.646; Fig. 3d, Supplementary Table 2; richness: see Supplementary Fig. 3b, d, Supplementary Table 2). However, the differential abundance analysis showed that the treatments within the laboratory experiment, i.e. the microbiota-environment combinations significantly affected the abundance of a large proportion of bacterial and fungal ASVs (bacteria: Fig. 3b, c; fungi: Fig. 3e, f). For both sun and shade microbiota, the abundance of specific ASVs differed between matching and mismatching conditions. Under sun matching conditions both bacteria and fungi showed a higher proportion (11.5% and 16.1%, respectively) of differential abundant taxa compared to mismatching conditions (6.3% and 3.2%, respectively). Under shade conditions only fungi showed a slightly higher proportion (20%) of differential abundant ASVs in matching conditions compared to mismatching conditions (17.1%) (Fig. 3b, c, e, f). Detailed taxonomic information is provided in the Supplementary Table 3a, b. For further contrasts see Supplementary Fig. 8.

**Fig. 3.**
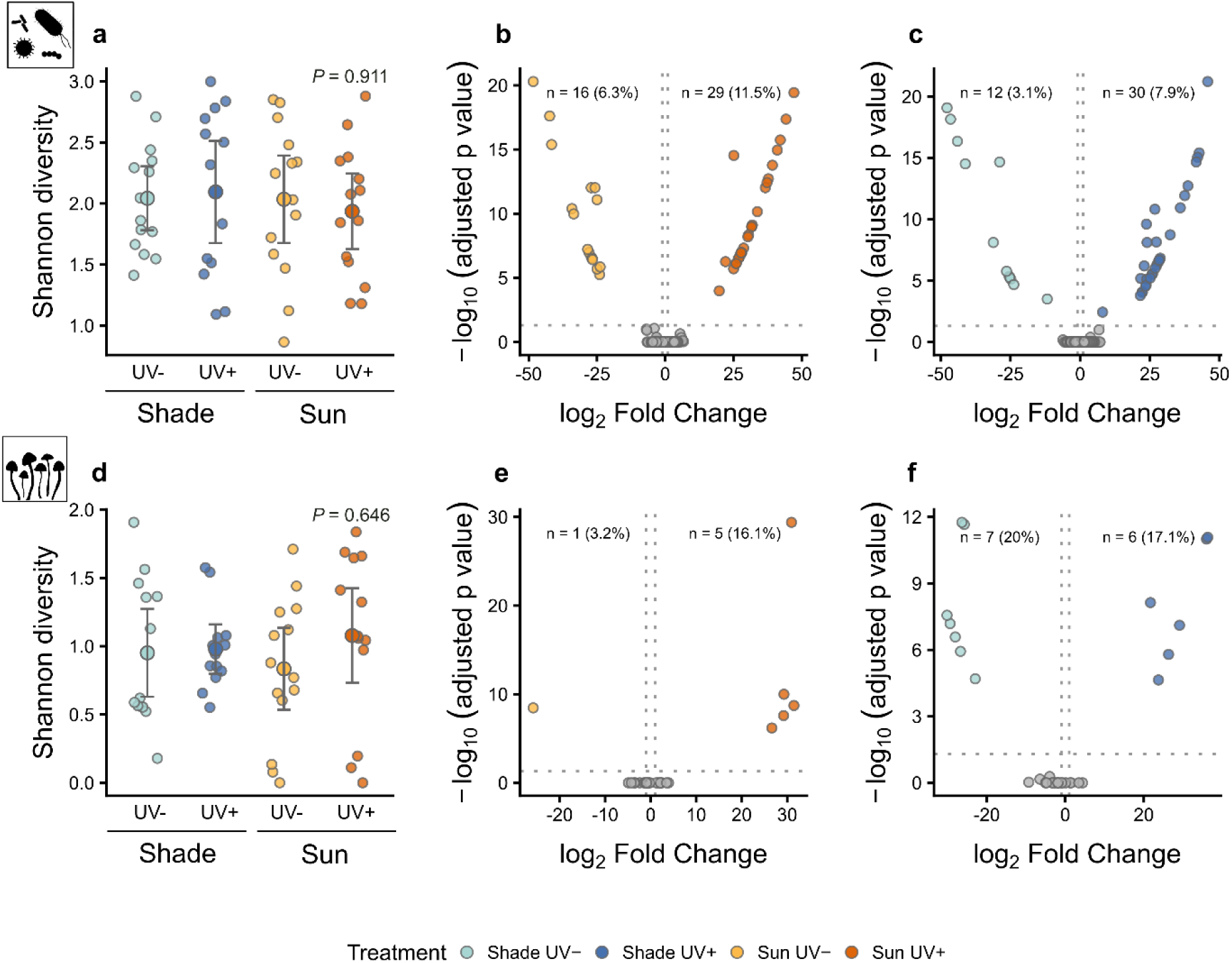
Alpha diversity and differential abundance of bacterial and fungal communities of treated oak ramets within an inoculation experiment. **a**, **c**, Shannon diversity index (*H’*) of bacterial (**a**) and fungal (**c**) leaf-associated communities across experimental treatments (*n* = 14 per treatment). Microbiota was extracted from leaves of treated oak ramets seven weeks after inoculation and start of UV exposure and analyzed via 16S rRNA gene (**a**) and ITS (**b**) amplicon sequencing. Large central circles and error bars indicate mean and 95% confidence intervals (CI), respectively. **b, c, e, f** Volcano plots illustrating differential abundance of individual bacteria (**b**, **c**) and fungal (**e**, **f**) amplicon sequence variants (ASVs) calculated using *DESeq2*. Separate contrasts were analyzed for sun (**b**, **e**) and shade (**c**, **f**) inoculated plants. Points represent individual ASVs plotted by their log_2_ fold change and – log_10_-transformed adjusted *P* values (*P_adj_*). Extreme log_2_ fold change values (±50) indicate ASVs with zero or near-zero values in one treatment, resulting in numerical capping by *DESeq2* (Love, Huber and Anders, 2014). Dotted lines indicate the significance thresholds (*P_adj_* < 0.05 and log_2_ fold change | > 1). Enriched ASVs are color-coded by experimental treatment. Icons were created by the authors or acquired and adapted from Phylopic.org (artists: F. Vaux, L. Simons). Results of the analyses of variance (ANOVA) are provided in the main text and Supplementary Table 2. For detailed taxonomic information of the significant bacterial and fungal ASVs see Supplementary Table 3a and b, respectively.

Plant performance was positively affected by matching microbiota-environment combinations as indicated by a significant interaction effect of inoculation type (origin of microbial inoculum) and UV exposure (ANOVA: NDVI: *F*_1,52_ = 5.691, *P* = 0.021; growth: *F*_1,52_ = 6.229, *P* = 0.016; Fig. 4, Supplementary Table 4a). This indicates that the microbial communities used for inoculation affected plant performance dependent on the experimental treatment, which means that matching microbiota-environment combinations affected plants differently than mismatching combinations: Plants inoculated with shade microbiota showed on average a 9.4% lower NDVI under UV treatment and plants inoculated with sun microbiota showed a 5.5% lower NDVI without UV treatment compared to those under respective matching conditions. Similarly, plants inoculated with shade microbiota showed a growth reduction of on average 42.3% under UV treatment and plants inoculated with sun microbiota showed a growth reduction of 21.5% without UV treatment compared to those under respective matching conditions. Neither the inoculation type (ANOVA: NDVI: *F*_1,52_ = 0.022, *P* = 0.882; growth: *F*_1,52_ = 1.003, *P* = 0.321) nor the experimental environment (UV exposure) (ANOVA: NDVI: *F*_1,52_ = 0.405, *P* = 0.527; growth: *F*_1,52_ = 0.615, *P* = 0.437) in isolation affected plant performance, which was also reflected in no significant contrast between the four experimental treatments (Tukey’s HSD test, Supplementary Table 4b). These findings support the notion that matching microbiota-environment combinations are required to promote plant performance, too.

**Fig. 4.**
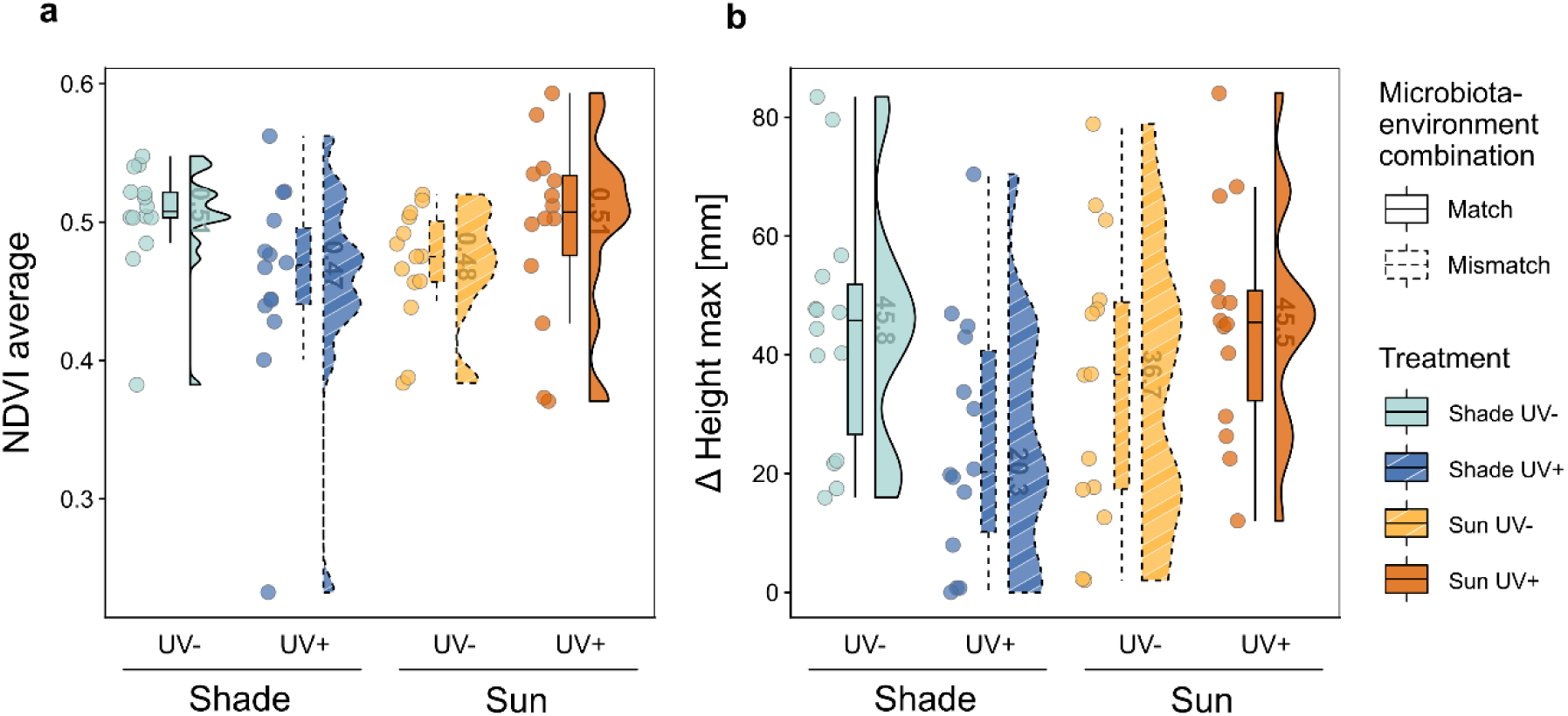
Plant phenotypic responses to microbial inoculation and environmental treatments. **a**, **b**, Performance of *Q. robur* ramets quantified by the Normalized Difference Vegetation Index (NDVI) (**a**) and difference in maximum height (ΔH_max)_ (**b**) after seven weeks of exposure to matched or mismatched microbiota-environment combinations. Data are presented as raincloud plots consisting of individual data points (*n* = 14 per treatment), boxplots (central horizontal line indicates median, box limits represent upper and lower quartiles, whiskers extend to 1.5 times the interquartile range (IQR)), and split-violin probability density distributions, with medians numerically annotated. Treatments are color-coded with additional indication for matching conditions by solid fills and mismatching conditions by striped patterns. Results of the analyses of variance (ANOVA) are provided in the main text and Supplementary Table 4.

### Network analysis

We used a correlation-based network analysis to gain a deeper understanding of interactions among bacteria, fungi, and the host plant. For both matching and mismatching microbiota-environment combinations we conducted separate analyses (Fig. 5a, b). The network representing plant-microbe interactions under matching conditions was more connected, less modular and exhibited higher average strength of associations compared to the network of mismatching condition (match: density = 0.172, modularity = 0.409, components = 1, average strength = 2.86; mismatch: density = 0.105, modularity = 0.550, components = 4, average strength = 2.38). Taxonomic information on microbial ASVs which were significantly correlated with plant traits are available in Supplementary Table 5.

**Fig. 5.**
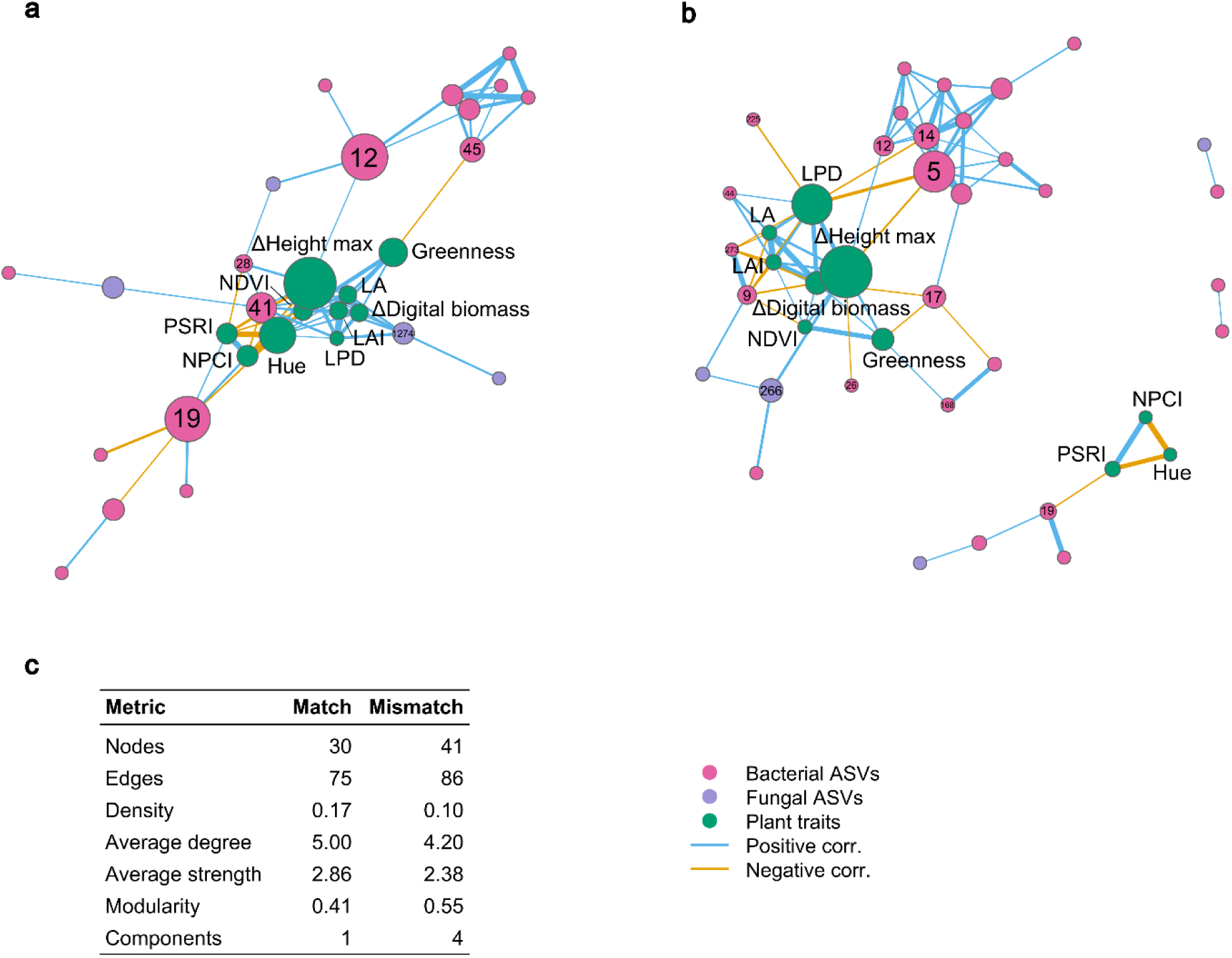
Correlation-based networks of microorganisms and plants traits under matching and mismatching conditions. **a**, **b**, Correlation-based networks showing interactions among bacterial amplicon sequence variants (ASVs), fungal ASVs and phenotypic plant traits under matching (**a**) and mismatching (**b**) microbiota-environment combinations. Nodes represent individual bacterial ASVs (pink), fungal ASVs (purple), or plant traits (green). Node sizes are proportional to their normalized betweenness centrality values. Edge colors indicate significant pairwise Spearman rank-sum correlations (*P* < 0.05), requiring co-occurrence in ≥ 9 samples), with blue and orange lines representing positive and negative correlations, respectively; edge width reflects correlation strength. ASVs which were significantly correlated with a plant trait are labelled (for detailed taxonomic information see Supplementary Table 5). **c**, Comparative summary of global network metrics for both matching and mismatching condition. Abbreviations: LA, Leaf area; LAI, Leaf area index; LPD, Light penetration depth; NDVI, Normalized Difference Vegetation Index; NPCI, Normalized Pigments Chlorophyll Ratio Index; PSRI, Plant Senescence Reflectance Index.

## Discussion

Our study indicates that intra-individual variation in leaf microbiota within tree crowns does not merely reflect microclimatic heterogeneity, but is also functionally relevant for the plant host. Microbiota-environment matching, i.e. the microbiota originating from an environment that was matched by the experimental condition, turned out to be a key determinant of plant performance. While the microbiota of sun-exposed leaves enhanced plant performance under UV exposure, equivalent beneficial effects were found when plants inoculated with shade microbiota were not exposed to UV in the laboratory experiment. These results emphasize that environment-specific microbiota occurring along the environmental gradient within tree crowns positively contribute to plant performance and underscore the functional relevance and ecological consequences arising from intra-individual variability in leaf-associated microbiota.

Studies at the individual host (within species) level have identified organ-specific microbiota, indicating strong niche differentiation and distinct functional potential (Arnold *et al*., 2025; Guo *et al*., 2025). Within plant organs, i.e. leaves, canopy position dependent differences in associated microbiota have been reported, consistent with our findings (Leff *et al*., 2015; Laforest-Lapointe, Messier and Kembel, 2016b; Herrmann *et al*., 2021; Addison *et al*., 2023). However, so far we lacked experimental evidence linking intra-individual variation in leaf microbiota to plant functioning. Using a mesocosm approach on oak ramets, we were able to directly link distinct microbiota to plant performance under defined environmental conditions. This way, we experimentally showed the relevance of *stratum*-specific leaf microbiota for host functioning. Previously, Herrera, (2017) described that intra-individual variation in functional plant traits can influence interactions with mutualists and antagonists, thereby affecting ecological networks and consequently plant health and function. We expand this concept to the plant-associated microbiota by demonstrating that within-host variability in leaf microbiota not only mirrors microhabitat heterogeneity but implies ecological consequences such as altered host performance.

The mechanisms underlying within-tree differentiation in leaf microbiota are likely multifactorial. Besides directly affecting microorganisms, environmental factors can also indirectly influence microbial communities through host-mediated changes in leaf traits (Bulgarelli *et al*., 2013; De Mandal and Jeon, 2023; Ma *et al*., 2026). In agreement with this effect, we measured light availability and temperature across canopy *strata* and found pronounced microclimatic variation consistent with other studies (De Frenne *et al*., 2021; Murakami *et al*., 2022). Further, canopy position was associated with differences in functional and physiochemical leaf traits, as demonstrated in another study on the same *Q. robur* individuals sampled in this study (Kong *et al*., 2025). Given this covariation in environmental conditions, host leaf traits and microbial communities along canopy *strata*, we propose that, rather than resulting from simple linear relationships, *stratum*-specific microbiota are shaped by interconnected interactions among environmental conditions, host responses and microorganisms.

The matching between environmental conditions and microbiota adapted to these conditions was key in explaining variation in plant performance among the inoculated oak ramets. Plants clearly benefited from microbial inoculation whenever the UV exposure at the origin of the inoculum matched the experimental UV radiation. This finding is consistent with previous studies on species level showing benefits of microbiota preadapted to prevailing environments (Carrell *et al*., 2022; Allsup, George and Lankau, 2023). In accordance with these studies, we emphasize that each plant-environment combination harbors environment-specific microbiota that provide the functional capacities needed under given conditions (Laforest-Lapointe, Messier and Kembel, 2016a; Lajoie, Maglione and Kembel, 2020). We highlight that this concept also applies to intra-individual environmental heterogeneity, i.e. to leaves at different positions within a tree canopy. Our results show no significant difference in alpha diversity for the treated plants of the inoculation experiment, while for each laboratory treatment we found certain differentially abundant microbial ASVs. This suggests that microbiome-mediated effects may depend more on the differential abundance and functional potential of particular community members than on microbial diversity per se (Trivedi *et al*., 2020; Moore *et al*., 2023; Poupin and González, 2024). Accordingly, one possible explanation for the benefit of environmental matching could be that matching conditions enrich microorganisms which enhance performance under these environmental conditions for example by growth promotion or pathogen suppression, while these functions might be reduced or absent under mismatching conditions (Hassani, Durán and Hacquard, 2018; Compant *et al*., 2019; De Mandal and Jeon, 2023). Alternatively, pathogen loads may be reduced by matching microbiota-environment conditions (Hassani, Durán and Hacquard, 2018; Ehau-Taumaunu and Hockett, 2023). However, these potential mechanisms remain speculative, as 16S rRNA gene and ITS amplicon sequencing does not allow for reliable inference on functions of specific taxa without complementary meta-omics or experimental validation (Knight *et al*., 2018; Matchado *et al*., 2024).

The advantageous effects of matching microbiota-environment combinations for the plant host were further reflected in the correlation-based network analysis. Although correlation-based networks do not necessarily reflect direct ecological interactions, but rather infer association based on co-variation (Faust and Raes, 2012; Poudel *et al*., 2016), network topology can help to reveal organizational patterns within biological systems. Here, the topology of microbial networks including plant traits and microbial taxa showed that communities under matching conditions were characterized by a single module and strong associations, indicating a highly integrated, stable system (Luo *et al*., 2023; Kajihara *et al*., 2025; Sun *et al*., 2025). In contrast, under mismatching conditions the network was more modular and the average association strength was reduced, similar to previous studies showing higher modularity or disrupted associations in stressful environments (de Vries *et al*., 2018; Yuan *et al*., 2021; Gao *et al*., 2022), but see (Hernandez *et al*., 2021; Luo *et al*., 2023; Kajihara and Hynson, 2024). We argue that in our study greater compartmentalization and larger network size may reflect an adaptive reorganization of the plant-microbe system in response to unfavorable conditions (Yuan *et al*., 2021; Cornell *et al*., 2023), whereas a higher average association strength within a single module network may indicate a more stable, functionally coherent system in non-stress environments. Moreover, the central positioning of plant traits within the matching network suggests that these traits act as key mediators among microbial groups contributing to network cohesion and stability (Faust and Raes, 2012; Toju, Tanabe and Sato, 2018; Banerjee *et al*., 2019). From a broader perspective, our results align with the concept of ecosystem coupling, where tightly linked systemic components support coordinated ecosystem functioning, while increasing stress can disrupt these interactions and undermine ecosystem functioning (Ochoa-Hueso *et al*., 2021).

Notably, studies in the context of environmental matching mainly focused on stress-related plant performance and further emphasized the role of rapid modulation of associated microbiota in plant acclimation under dynamic or changing environments (Carrell *et al*., 2022; Allsup, George and Lankau, 2023; Zieschank *et al*., 2025). As rapid anthropogenic changes often outpace natural plant adaptation (Lau and Lennon, 2012; Trivedi *et al*., 2022), we agree that microbial assistance may represent a promising mechanism to buffer limitations in plant responses to environmental stress. However, beyond the application of technological interventions (as used in transplantation experiments), our results highlight the broad benefits of natural and functionally diverse microbial communities in biological systems (Junker and Farwig, 2025). By revealing the significance of intra-individual microbial variability, we emphasize the need to ensure high levels of microbial diversity as a prerequisite for plant-environment specific microbiota and conclude that efforts in ecosystem conservation should implement the protection of microorganisms. Therefore, our data strengthen the case for microbial conservation, which ultimately aims to protect intact, functional natural systems (Junker and Farwig, 2025). With this study, we broaden current understanding of microbiome contributions to plant functioning and health by demonstrating that *stratum*-specific variation in leaf microbiota within individual trees is not solely a reflection of environmental heterogeneity, but represents a functionally relevant feature through which locally adapted microbiota collectively support plant performance.

## Acknowledgements

We thank Marco Göttig for cultivating the plant material. Further, we thank Esther Meißner, Sebastian Achilles and Mona Schreiber for their help during field work as well as Vicki Tough, Kathina Müßig, Lara Lohmann, and Alex Stylianou for supporting leaf collection as professional climbers. For essential support in constructing the UV-B LED modules, we thank the people of the University workshop. We thank Manuel Pitzer for assistance in laboratory work, technical support and the valuable input and discussions on the topic of the manuscript. We thank Lisa Bald and Kristian Peters for support with data management and Doreen Meier as project coordinator.

## Funding Statement

This work was funded by the LOEWE research initiative of the State of Hesse, Germany, through the Ministry of Science and Arts (HMWK), as part of the LOEWE research cluster Tree-M (LOEWE/2/15/519/03/08.001(0002)/88 and by the Deutsche Forschungsgemeinschaft (DFG, German Research Foundation, JU 2856/10-1 and OP 219/20-1) in the framework of RU 5571 PhytOakmeter.

## Author Contributions

R.R.J., H.A. and L.S., designed the study, R.R.J., H.A., L.S., A.R.W., A.B., N.F., S.P., L.O., C.L., J.B. contributed to the conception of the work, L.S., A.R.W., A.N.A.R., S.W., T.T. and F.K. conducted the field work. L.O. provided the plant material for the laboratory experiment. Analysis was carried out by L.S., R.R.J. and H.A. with support from S.J. and A.L. The original draft was written by L.S. and R.R.J. All authors reviewed and edited the final manuscript.

## Competing interests

The authors declare no competing interests.

## Data availability

Raw sequencing data have been deposited at the European Nucleotide Archive (ENA; accession number: ERP196146). All data and the R script used for data analysis will be made publicly available via the dataPLANT publication ARChive by nfdi4plants upon publication of the manuscript.

## Notes

### Competing Interest Statement

The authors have declared no competing interest.

