## Supplementary Material for "Intra-individual variation in leaf microbiota matches within-crown environmental heterogeneity and promotes tree performance"

#### Supplementary Figures

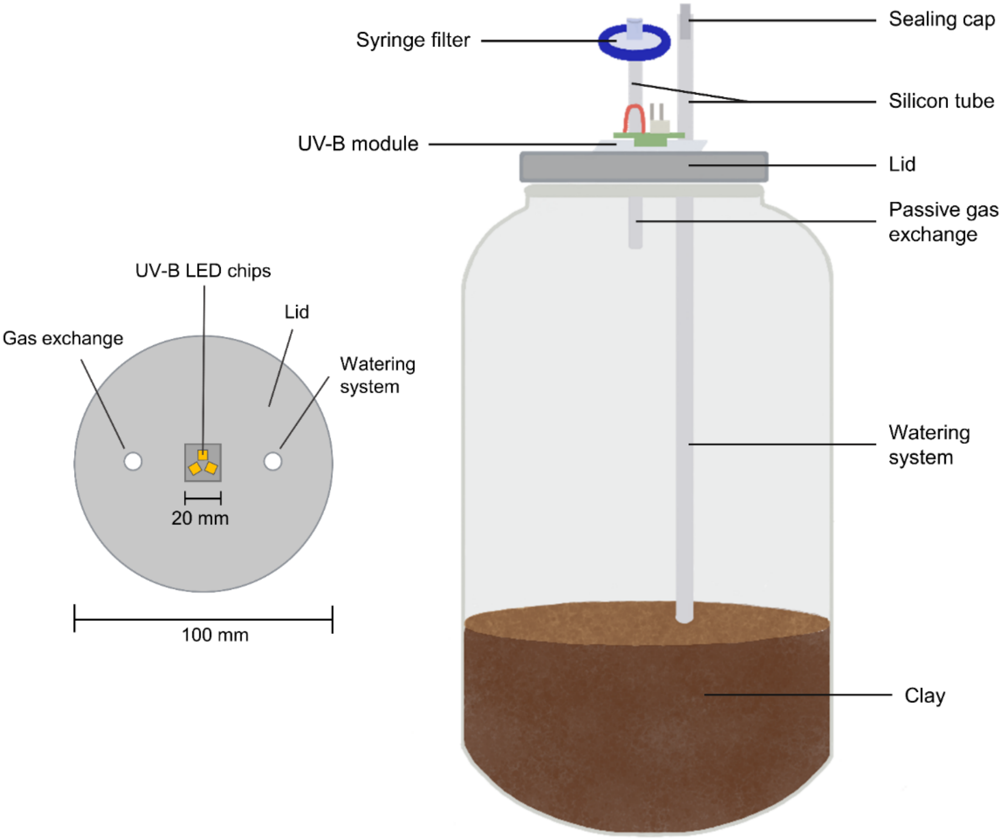


**Supplementary Fig 1. Mesocosm setup and detailed view of the UV-B module**. A mesocosm consists of a 4250 mL glass jar paired with the appropriate, sterilizable lid. The lid is attached to a watering system consisting of a 4 x 7 mm silicon tube inserted into the sterilized substrate at the one end and closed with a 4 mm sealing cap at the other end. A syringe filter (SFCA, 25 mm diam., 0.2 µm) attached to a shorter 4 x 7 mm silicon tube allows for passive gas exchange. Three narrowband LED chips emitting at 310- 315 nm are installed inside the lid. A broad viewing angle of 120° and a flat quartz glass window ensure Lambertian emission. UV intensity can be adjusted continuously from 0.007 to 0.021 mW/cm$^2$.

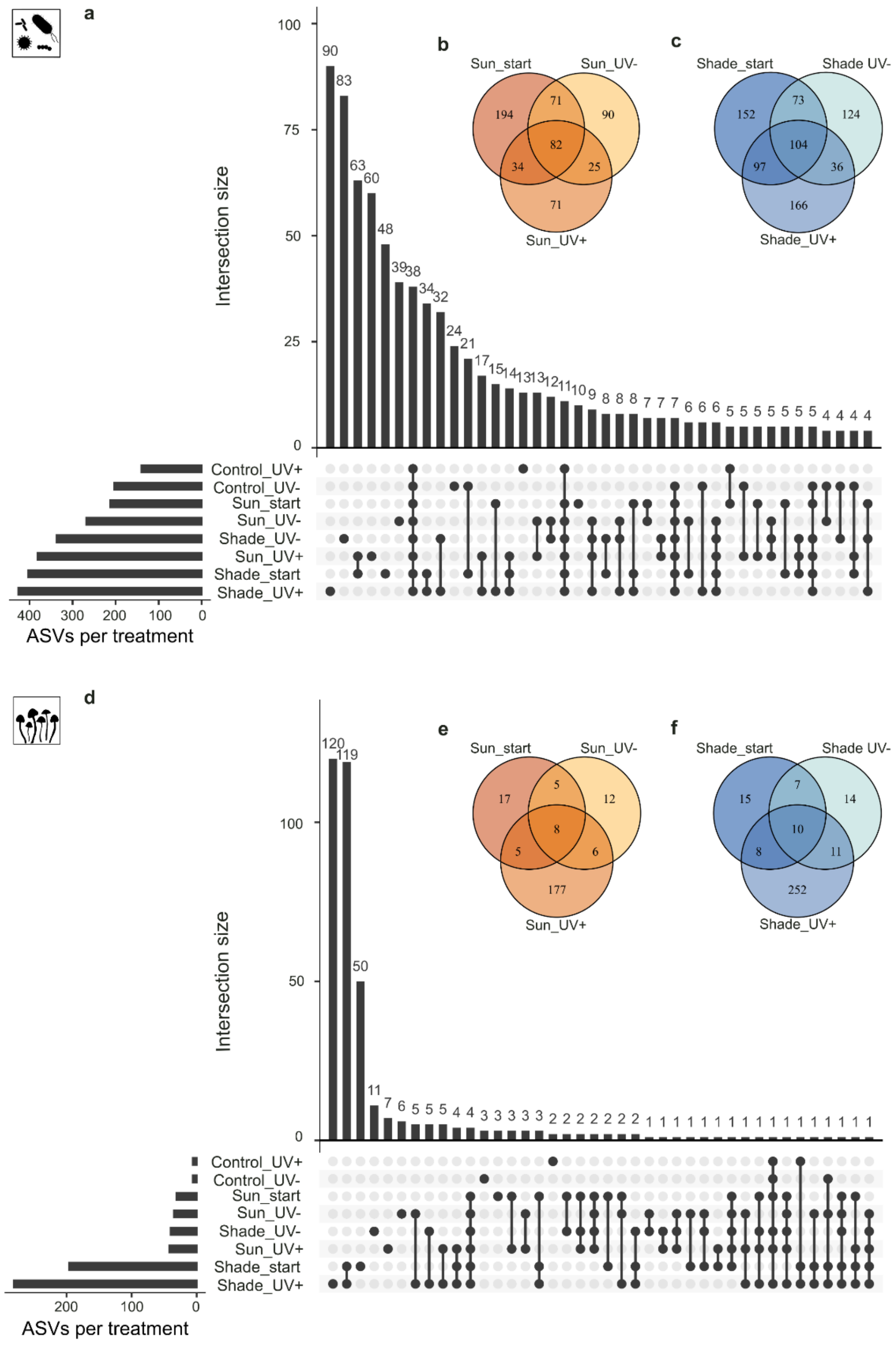


**Supplementary Fig. 2 Shared and unique amplicon sequence variants (ASVs) across experimental groups**. **a-c**, Bacterial ASV distribution. **a**, Upset plot displaying the distinct intersections of bacterial ASVs among the eight experimental groups. **b**, **c**, Venn diagrams showing the overlapping bacterial ASVs between initial field samples (start) used as microbial inocula and treated oak ramets after seven weeks for sun (**b**) and shade (**c**) groups. **d-f**, Fungal ASV distribution, showing the corresponding Upset plot (**d**) and Venn diagrams for sun (**e**) and shade (**f**) groups. Icons were created by the authors or acquired and adapted from Phylopic.org (artists: F. Vaux, L. Simons).

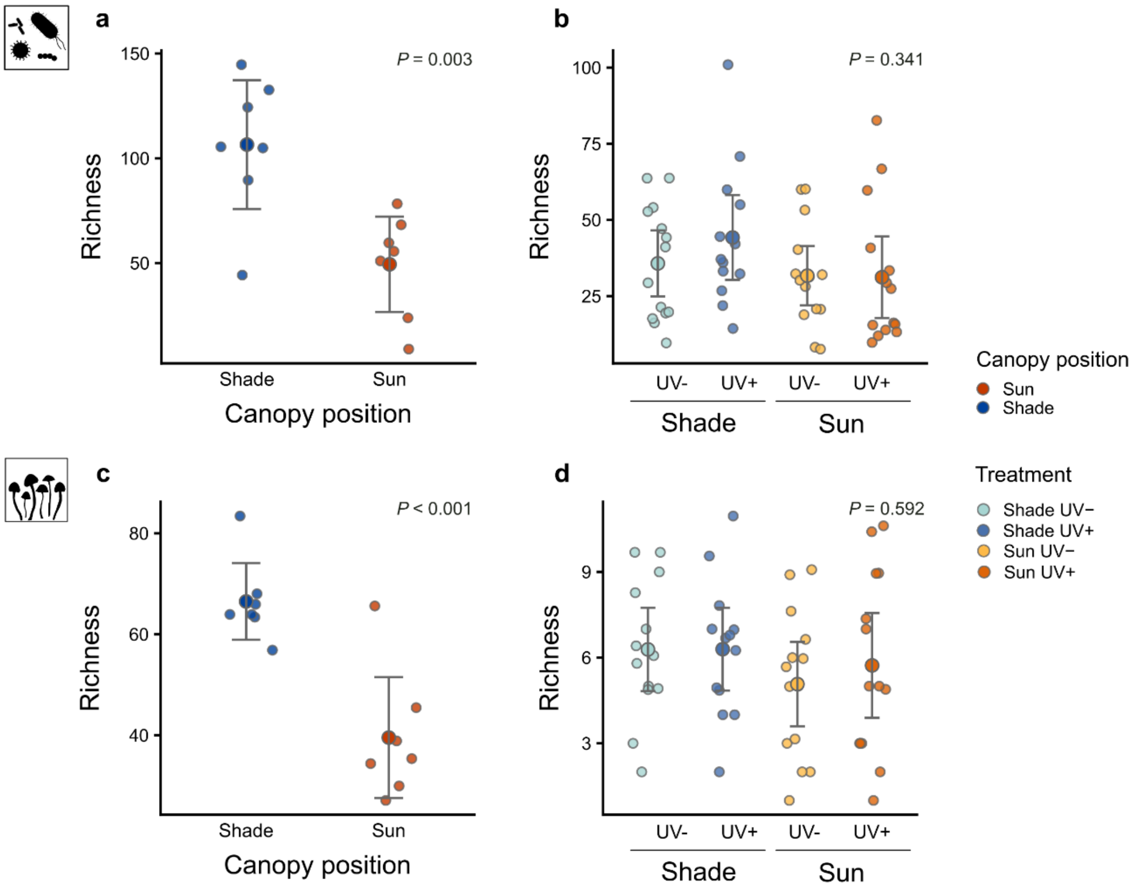


**Supplementary Fig. 3 Microbial species richness across canopy positions and laboratory treatments**. **a**, **c**, Species richness of bacterial (**a**) and fungal (**c**) communities sampled from different canopy positions (sun, orange; shade, blue) of *n* = 7 *Quercus robur* forest trees. Large central circles and error bars indicate the group mean and 95% confidence intervals, respectively. **b**, **d**, Species richness of bacterial (**b**) and fungal (**d**) communities across the four laboratory treatments of the inoculation experiment after 7 weeks (visualized as described in **a**, **c**). Richness was calculated from rarified 16S rRNA gene (**a**, **c**) and ITS (**c**, **d**) amplicon sequencing data. Results of the analyses of variance (ANOVA) are provided in Supplementary Table 2. Icons were created by the authors or acquired and adapted from Phylopic.org (artists: F. Vaux, L. Simons).

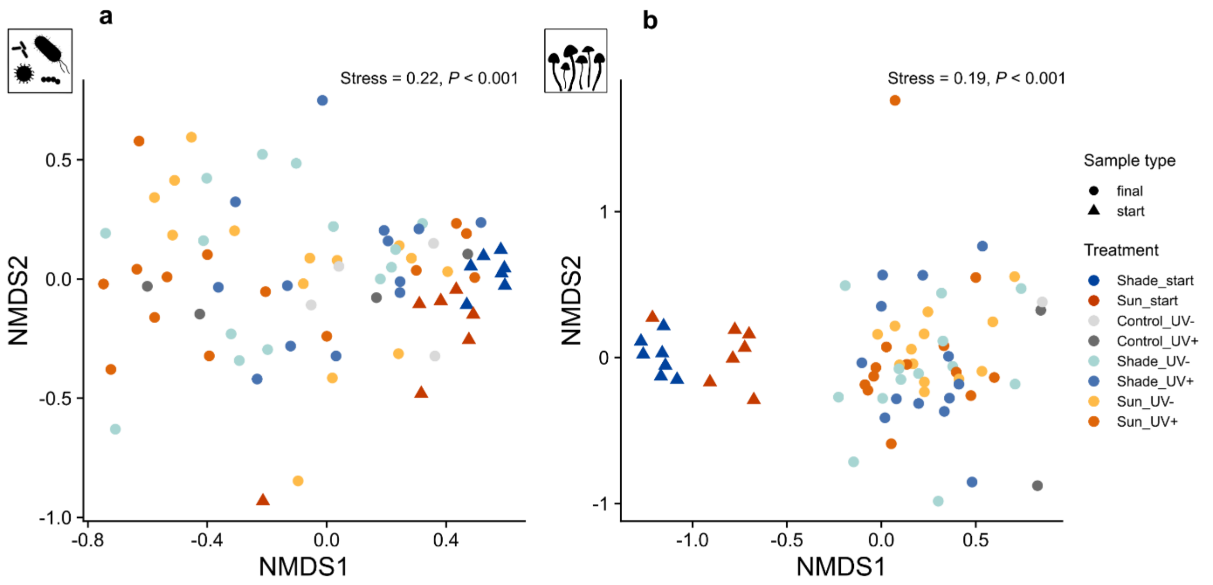


**Supplementary Fig. 4 Bacterial and fungal community composition across canopy positions and experimental treatments. a**, **b** Non-metric multidimensional scaling (NMDS) ordinations of bacterial (**a**) and fungal (**b**) community composition based on Bray-Curtis distances. Microbial count data from 16S rRNA gene (**a**) and ITS (**b**) amplicon sequencing were normalized using cumulative sum scaling (CSS), followed by min-max normalization. Individual data points represent biological replicates, shaped by sample type (triangles for microbiota of field samples; circles for microbiota of laboratory samples) and color-coded by treatment group. Icons were created by the authors or acquired and adapted from Phylopic.org (artists: F. Vaux, L. Simons). Global goodness-of-fit for each ordination is indicated by the annotated stress value. Differences in community structure across treatments were statistically evaluated using PERMANOVA.

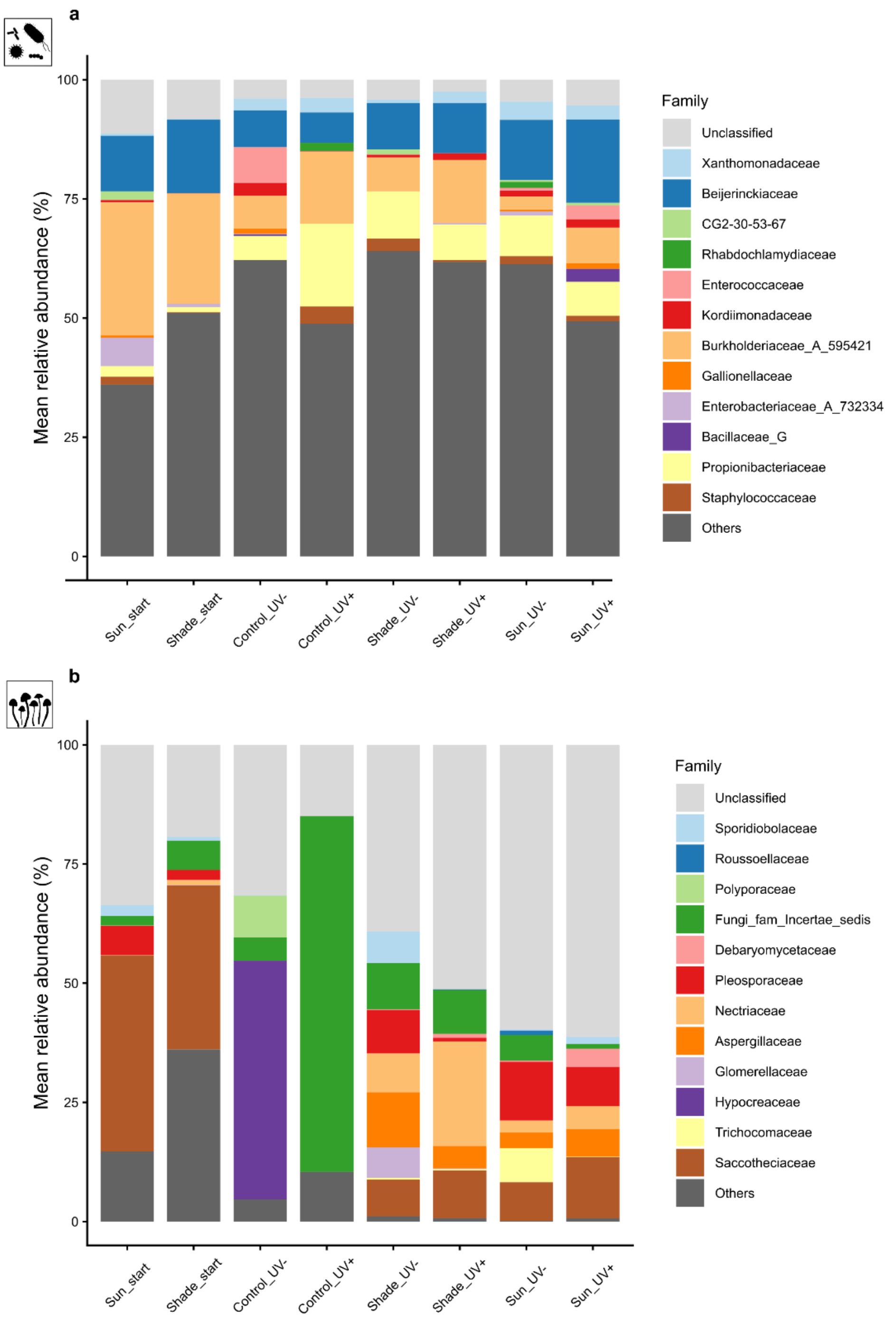


**Supplementary Fig. 5 Relative abundance of bacterial and fungal families across experimental groups. a**, **b**, Stacked barplots representing the mean relative abundance of bacterial (**a**) and fungal (**b**) families across the eight experimental groups. Bars display the top 12 most abundant families across all samples; unclassified taxa at the family level are indicated in light grey, while dark grey summarizes all remaining less-abundant taxa clustered as "Others". Treatment groups are displayed on the x-axis in chronological order ranging from initial field samples (start) used as microbial inocula to samples of treated oak ramets after seven weeks of experimental conditions (control, sun, shade inoculated plants under UV+ and UV- environments). Icons were created by the authors or acquired and adapted from Phylopic.org (artists: F. Vaux, L. Simons).

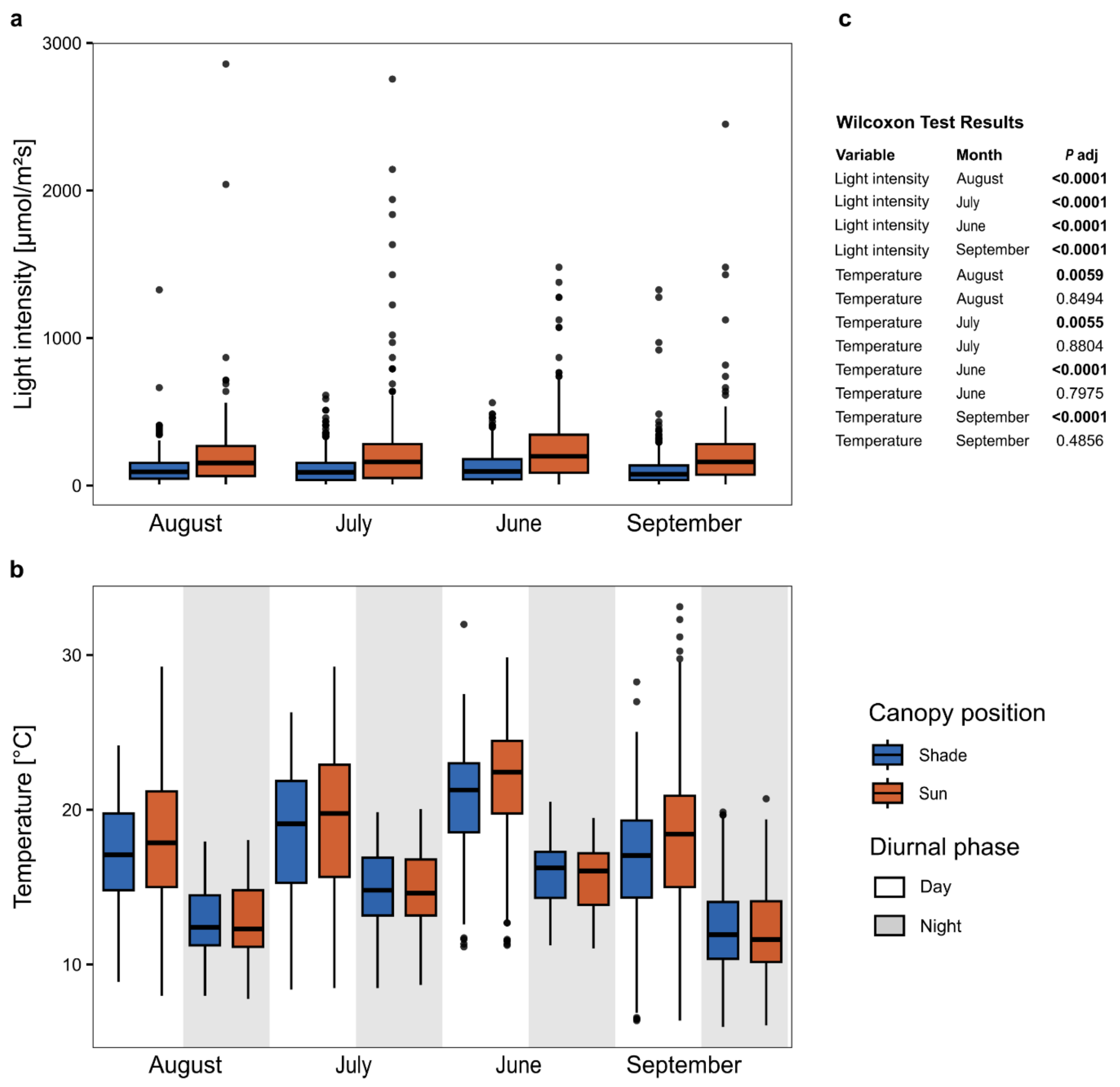


**Wilcoxon Test Results**

| Variable | Month | *P* adj |
|---|---|---|
| Light intensity | August | **<0.0001** |
| Light intensity | July | **<0.0001** |
| Light intensity | June | **<0.0001** |
| Light intensity | September | **<0.0001** |
| Temperature | August | **0.0059** |
| Temperature | August | 0.8494 |
| Temperature | July | **0.0055** |
| Temperature | July | 0.8804 |
| Temperature | June | **<0.0001** |
| Temperature | June | 0.7975 |
| Temperature | September | **<0.0001** |
| Temperature | September | 0.4856 |

**Supplementary Fig. 6 Seasonal and diurnal variation in microclimatic conditions along canopy *strata*. a**, Light intensity at the two canopy positions sun and shade (referring to daytime hours only) across seasons (July to September). **b**, Diurnal temperature at both canopy positions across seasons. Grey shaded background indicates night. For each boxplot, the center horizontal line represents the median, the box limits show the upper and lower quartiles, whiskers extend to 1.5 times the interquartile range (IQR), and points show outliers. The colors are indicative for canopy position (sun, orange; shade, blue). **c**, Pairwise comparisons of the tested environmental variables between canopy positions sun and shade. Values represent adjusted *P* values ($P_{adj}$) calculated via Wilcoxon rank-sum tests, with statistically significant differences ($P_{adj}$ < 0.001) indicated in bold. Data were collected from permanently installed dataloggers at both canopy positions in each sampled *Quercus robur* tree.

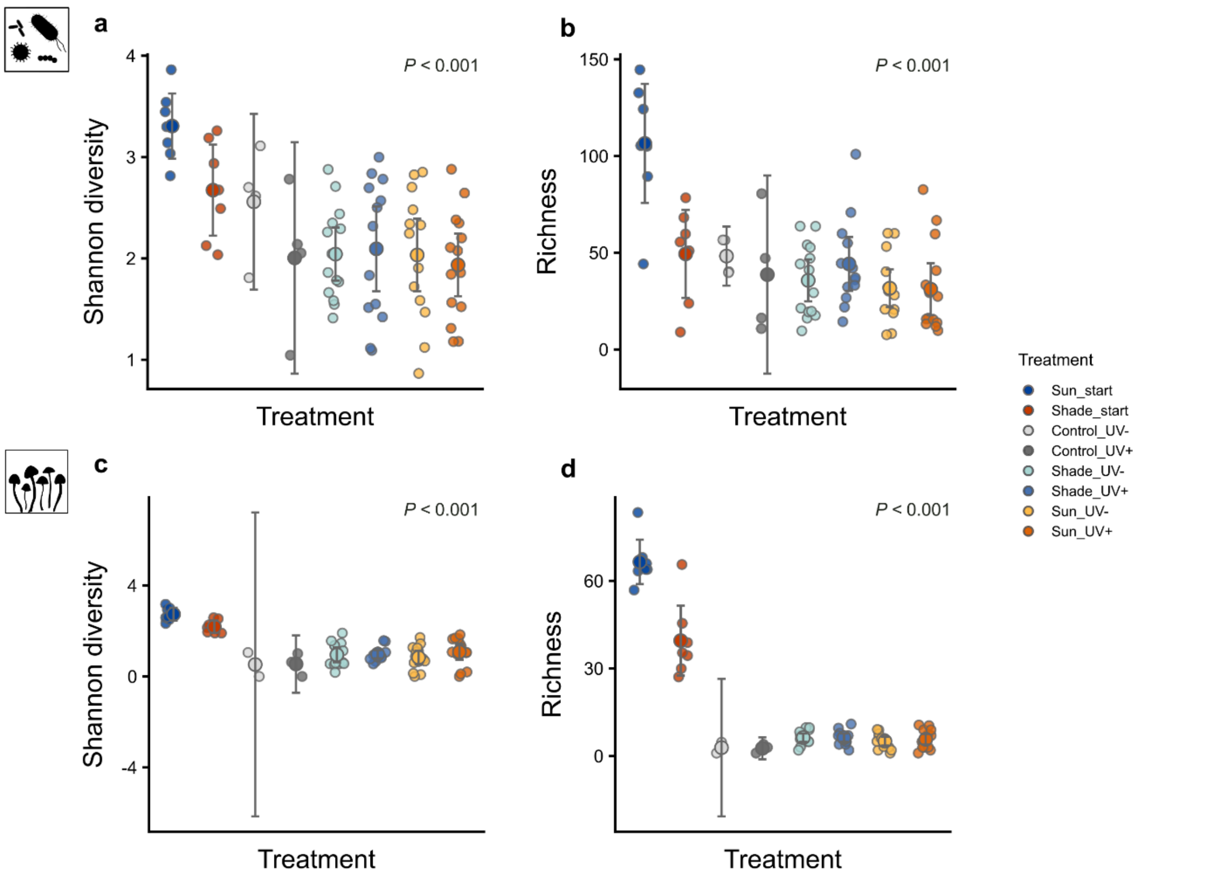


**Supplementary Fig. 7 Comprehensive bacterial and fungal alpha diversity across field samples and laboratory treatments. a**, **b**, Alpha diversity of bacterial communities across all experimental groups, quantified by the Shannon diversity index (*H′*) (**a**) and species richness (**b**). Groups include field samples (used as microbial inocula) and the laboratory experimental treatments, including controls. Microbial communities were sampled from *n* = 7 *Quercus robur* trees and *n* = 14 treated oak ramets (*n* = 4 for controls) and characterized via 16S rRNA gene amplicon sequencing (for alpha diversity calculation rarified data were used). Large central circles and error bar indicate the group mean and the 95% confidence intervals, respectively. **c**, **d** Alpha diversity of fungal communities across the identical field and laboratory groups described above, quantified by Shannon diversity (*H'*) (**c**) and species richness (**d**) based on rarified ITS amplicon sequencing data (visualized as described in **a**, **b**). Icons were created by the authors or acquired and adapted from Phylopic.org (artists: F. Vaux, L. Simons). Results of the analyses of variance are provided in Supplementary Table 2.

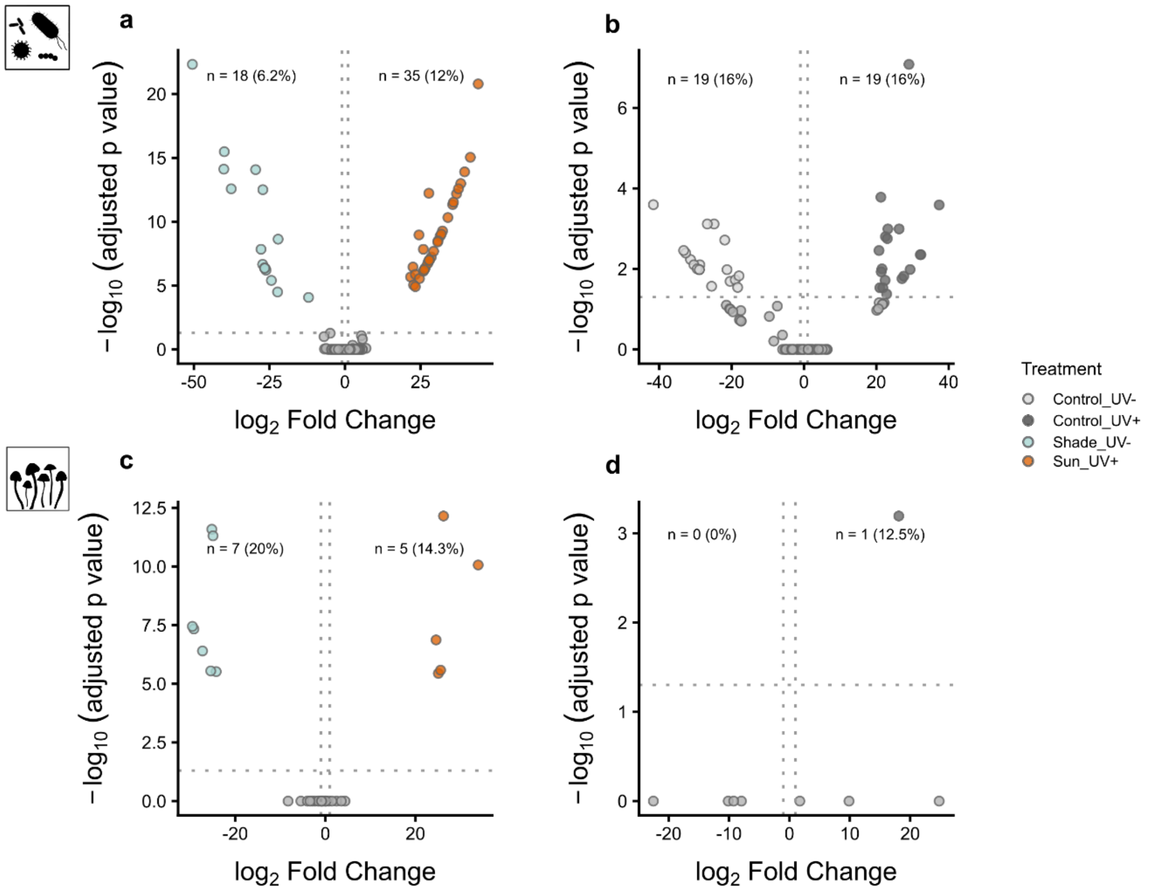


**Supplementary Fig. 8 Additional differential abundance contrasts of bacterial and fungal communities. a–d**, Volcano plots illustrating the differential abundance of individual bacterial (**a**, **b**) and fungal (**c**, **d**) amplicon sequence variants (ASVs) calculated using *DESeq2* for specific treatment contrasts. Pairwise comparisons were performed to evaluate differences between the two matching microbiota–environment combinations (Shade UV- vs. Sun UV+) (**a**, **c**) and between the two uninoculated control groups (Control UV+ vs. Control UV-) (**b**, **d**). Points represent individual ASVs plotted by their $\log_2$ fold change and – $\log_{10}$-transformed adjusted *P* values ($P_{adj}$). Extreme $\log_2$ fold change values (±40) indicate ASVs with zero or near-zero values in one treatment, resulting in numerical capping by *DESeq2*[97]. Dotted lines indicate the significance thresholds ($P_{adj} < 0.05$ and $\log_2$ fold change | > 1). Enriched ASVs are color-coded by experimental treatment. Icons were created by the authors or acquired and adapted from Phylopic.org (artists: F. Vaux, L. Simons). For detailed taxonomic information of the significant bacterial and fungal ASVs see Supplementary Table 3a and b, respectively.

### Supplementary Tables

**Supplementary Table 1**. Experimental groups. **a**, Field samples according to the position within the tree canopy of *Quercus robur* (sun/ shade). **b**, Treatments of the inoculation experiment. Isolated microbiota from field samples, either of sun or shade leaves, were transferred to germ-reduced oak clones of the same species. According to the UV exposure in the laboratory, the conditions either matched or mismatched the UV condition of the environment of origin of the microbial inoculum (sun /shade). Colors correspond to Fig. 1, 3 and 4 of the main text.

a) Field samples

| Canopy position | Sample type | Treatment |
|---|---|---|
| Shade | start | Start_Sun |
| Sun | start | Start_Shade |

b) Inoculation experiment

| Inoculation type | Sample type | Experimental condition | Microbiota-environment combination | Treatment |
|---|---|---|---|---|
| Control | final | UV- | NA | Control_UV- |
| Control | final | UV+ | NA | Control_UV+ |
| Shade | final | UV- | Match | Shade_UV- |
| Shade | final | UV+ | Mismatch | Shade_UV+ |
| Sun | final | UV- | Mismatch | Sun_UV- |
| Sun | final | UV+ | Match | Shade_UV+ |

**Supplementary Table 2.** Results of a two-factorial ANOVA testing for differences in Shannon diversity and richness of leaf-associated microbiota across canopy *strata* (field) and experimental treatments (experiment), using canopy position (sun/ shade) and treatment as explanatory variables, respectively. Significant results are indicated in bold.

| **Sample group** | **Diversity metric** | **Factor** | **Bacteria** | | **Fungi** | | **Figure** |
|---|---|---|---|---|---|---|---|
| | | | $F_{1,12}$ | *P* | $F_{1,12}$ | *P* | |
| Field | Shannon diversity | Canopy position | 7.834 | **0.016** | 11.946 | **0.005** | Fig. 2 |
| | | | $F_{1,12}$ | *P* | $F_{1,12}$ | *P* | |
| Field | Richness | Canopy position | 13.36 | **0.003** | 21.65 | **<0.001** | Supplementary Fig. 2 |
| | | | $F_{3,51}$ | *P* | $F_{3,50}$ | *P* | |
| Experiment | Shannon diversity | Treatment | 0.178 | 0.911 | 0.556 | 0.646 | Fig. 3 |
| | | | $F_{3,51}$ | *P* | $F_{3,50}$ | *P* | |
| Experiment | Richness | Treatment | 1.143 | 0.341 | 0.642 | 0.592 | Supplementary Fig. 2 |
| | | | $F_{7,69}$ | *P* | $F_{7,65}$ | *P* | |
| Global | Shannon diversity | Treatment | 5.706 | **<0.001** | 18.210 | **<0.001** | Supplementary Fig. 6 |
| | | | $F_{7,69}$ | *P* | $F_{3,50}$ | *P* | |
| Global | Richness | Treatment | 9.257 | **<0.001** | 0.642 | **<0.001** | Supplementary Fig. 6 |

**Supplementary Table 3a.** Differentially abundant bacterial ASVs across treatments (shown in Fig. 2, 3 of the main text). Statistical parameters were calculated using *DESeq2* .

| Contrast | Direction | Base Mean | Log2 Fold Change | *P* | *Padj* | ASV | Domain | Phylum | Class | Order | Family | Genus | Species |
|---|---|---|---|---|---|---|---|---|---|---|---|---|---|
| Sun start vs. Shade start | Shade start | 143.741 | -25.879 | <0.0001 | <0.0001 | ASV32 | Bacteria | Actinomycetota | Actinomycetes | Actinomycetales | Kineococcaceae | Kineococcus | Kineococcus radiotolerans |
| Sun start vs. Shade start | Shade start | 73.054 | -24.368 | <0.0001 | 0.0003 | ASV21 | Bacteria | Pseudomonadota | Gammaproteobacteria | Enterobacterales_737866 | | | |
| Sun start vs. Shade start | Sun start | 22.221 | 26.303 | 0.0001 | 0.0125 | ASV154 | Bacteria | Actinomycetota | Actinomycetes | Actinomycetales | | | |
| Sun start vs. Shade start | Sun start | 51.022 | 21.649 | <0.0001 | 0.0125 | ASV228 | Bacteria | Pseudomonadota | Gammaproteobacteria | Burkholderiales | Burkholderiaceae_A_595421 | | |
| Sun start vs. Shade start | Shade start | 24.338 | -25.675 | 0.0002 | 0.0128 | ASV222 | Bacteria | Pseudomonadota | Gammaproteobacteria | Burkholderiales | Burkholderiaceae_A_595421 | | |
| Sun start vs. Shade start | Sun start | 113.399 | 23.643 | 0.0002 | 0.0128 | ASV332 | Bacteria | Pseudomonadota | Gammaproteobacteria | Burkholderiales | Burkholderiaceae_A_595421 | | |
| Sun start vs. Shade start | Sun start | 4.191 | 25.502 | 0.0002 | 0.0128 | ASV2012 | Bacteria | Pseudomonadota | Alphaproteobacteria | Sphingomonadales | Sphingomonadaceae_486827 | Sphingomonas_L_486704 | Sphingomonas_L_486704 mali |
| Sun start vs. Shade start | Sun start | 2.768 | 25.034 | 0.0002 | 0.0132 | ASV1075 | Bacteria | Bacteroidota | Bacteroidia | Flavobacteriales_B_877923 | Weeksellaceae | Kaistella | Kaistella sp000023725 |
| Sun start vs. Shade start | Sun start | 30.844 | 24.814 | 0.0003 | 0.0133 | ASV118 | Bacteria | Actinomycetota | Actinomycetes | Mycobacteriales | Mycobacteriaceae | Corynebacterium | |
| Sun start vs. Shade start | Shade start | 35.122 | -22.779 | 0.0004 | 0.0157 | ASV47 | Bacteria | Pseudomonadota | Alphaproteobacteria | Rhizobiales_505101 | Beijerinckiaceae | Methylobacterium | Methylobacterium sp001424705 |
| Sun start vs. Shade start | Shade start | 13.277 | -21.687 | 0.0005 | 0.0157 | ASV134 | Bacteria | Actinomycetota | Actinomycetes | Mycobacteriales | Micromonosporaceae | | |
| Sun start vs. Shade start | Shade start | 9.519 | -23.729 | 0.0005 | 0.0157 | ASV214 | Bacteria | Bacteroidota | Bacteroidia | Flavobacteriales_B_877923 | Weeksellaceae | | |
| Sun start vs. Shade start | Shade start | 3.047 | -24.027 | 0.0004 | 0.0157 | ASV1724 | Bacteria | Bacteroidota | Bacteroidia | Cytophagales_B | Hymenobacteraceae | Hymenobacter_910554 | Hymenobacter sp004684095 |
| Sun start vs. Shade start | Sun start | 4.741 | 23.811 | 0.0005 | 0.0157 | ASV457 | Bacteria | Pseudomonadota | Alphaproteobacteria | Acetobacterales | Acetobacteraceae | | |
| Sun start vs. Shade start | Sun start | 10.655 | 23.46 | 0.0006 | 0.017 | ASV520 | Bacteria | Actinomycetota | Actinomycetes | Actinomycetales | Brevibacteriaceae | Brevibacterium | |
| Sun start vs. Shade start | Shade start | 7.758 | -22.919 | 0.0008 | 0.0204 | ASV69 | Bacteria | Pseudomonadota | Alphaproteobacteria | Sphingomonadales | Sphingomonadaceae_486827 | Sphingomonas_L_486704 | Sphingomonas_L_486704 olei |
| Sun start vs. Shade start | Shade start | 5.876 | -22.89 | 0.0008 | 0.0204 | ASV277 | Bacteria | Pseudomonadota | Gammaproteobacteria | Burkholderiales | Burkholderiaceae_A_595421 | | |
| Sun start vs. Shade start | Shade start | 8.768 | -22.37 | 0.001 | 0.0218 | ASV75 | Bacteria | Bacteroidota | Bacteroidia | Cytophagales_B | Hymenobacteraceae | Hymenobacter_910554 | Hymenobacter setariae |
| Sun start vs. Shade start | Shade start | 66.071 | -22.361 | 0.001 | 0.0218 | ASV150 | Bacteria | Actinomycetota | Actinomycetes | Propionibacteriales | Nocardioidaceae | Nocardioides_A_392796 | |
| Sun start vs. Shade start | Shade start | 1.684 | -22.585 | 0.0009 | 0.0218 | ASV375 | Bacteria | Pseudomonadota | Alphaproteobacteria | Sphingomonadales | Sphingomonadaceae_486827 | Sphingomonas_L_486704 | |

| | | | | | | | | | | | | | |
|---|---|---|---|---|---|---|---|---|---|---|---|---|---|
| Sun start vs. Shade start | Shade start | 4.051 | -22.494 | 0.001 | 0.0218 | ASV495 | Bacteria | Bacillota_I | Bacilli_A | Paenibacillales | Paenibacillaceae_367444 | Saccharibacillus | Saccharibacillus sacchari |
| Sun start vs. Shade start | Shade start | 1.976 | -22.001 | 0.0012 | 0.0251 | ASV401 | Bacteria | Pseudomonadota | Alphaproteobacteria | Acetobacterales | Acetobacteraceae | | |
| Sun start vs. Shade start | Shade start | 20.003 | -21.812 | 0.0014 | 0.0253 | ASV172 | Bacteria | Bacillota_I | Bacilli_A | Lactobacillales | Streptococcaceae | Lactococcus_A_346120 | |
| Sun start vs. Shade start | Shade start | 32.122 | -21.662 | 0.0015 | 0.0253 | ASV213 | Bacteria | Actinomycetota | Actinomycetes | Propionibacteriales | Nocardioidaceae | Nocardioides_A_392796 | |
| Sun start vs. Shade start | Shade start | 2.064 | -21.703 | 0.0014 | 0.0253 | ASV329 | Bacteria | Bacteroidota | Bacteroidia | Flavobacteriales_B_877923 | Weeksellaceae | | |
| Sun start vs. Shade start | Shade start | 2.39 | -21.731 | 0.0014 | 0.0253 | ASV795 | Bacteria | Pseudomonadota | Alphaproteobacteria | Acetobacterales | Acetobacteraceae | | |
| Sun start vs. Shade start | Shade start | 3.112 | -21.372 | 0.0017 | 0.0281 | ASV206 | Bacteria | Actinomycetota | Actinomycetes | Actinomycetales | Dermatophilaceae | | |
| Sun start vs. Shade start | Shade start | 15.816 | -21.251 | 0.0018 | 0.0288 | ASV397 | Bacteria | Bacillota_I | Bacilli_A | Lactobacillales | Streptococcaceae | Streptococcus | |
| Sun start vs. Shade start | Shade start | 7.422 | -21.054 | 0.002 | 0.0307 | ASV188 | Bacteria | Actinomycetota | Actinomycetes | Mycobacteriales | Geodermatophilaceae | | |
| Sun start vs. Shade start | Shade start | 2.965 | -20.557 | 0.0025 | 0.0373 | ASV341 | Bacteria | Pseudomonadota | Alphaproteobacteria | | | | |
| Sun start vs. Shade start | Shade start | 2.695 | -20.518 | 0.0026 | 0.0373 | ASV379 | Bacteria | Acidobacteriota | Terriglobia | Bryobacterales | Bryobacteraceae | KBS-96 | KBS-96 sp000381625 |
| Sun start vs. Shade start | Shade start | 91.323 | -20.303 | 0.0029 | 0.0401 | ASV351 | Bacteria | Bacillota_I | Bacilli_A | Bacillales_B_306087 | DSM-18226_301387 | | |
| Sun start vs. Shade start | Shade start | 19.2 | -20.163 | 0.0031 | 0.0409 | ASV97 | Bacteria | Pseudomonadota | Gammaproteobacteria | Enterobacterales_737866 | Enterobacteriaceae_A_726016 | Gibbsiella | Gibbsiella greigii |
| Sun start vs. Shade start | Shade start | 20.423 | -20.137 | 0.0031 | 0.0409 | ASV252 | Bacteria | Pseudomonadota | Alphaproteobacteria | Rhodobacterales | Rhodobacteraceae | Paracoccus | |
| Sun start vs. Shade start | Shade start | 2.437 | -20.044 | 0.0032 | 0.0415 | ASV56 | Bacteria | Bacteroidota | Bacteroidia | Cytophagales_B | Hymenobacteraceae | Solirubrum | |
| Sun start vs. Shade start | Shade start | 4.481 | -19.914 | 0.0035 | 0.0429 | ASV93 | Bacteria | Pseudomonadota | Alphaproteobacteria | Rhizobiales_505101 | Rhizobiaceae | | |
| Sun start vs. Shade start | Shade start | 29.691 | -19.759 | 0.0037 | 0.045 | ASV270 | Bacteria | Pseudomonadota | Gammaproteobacteria | Pseudomonadales_A_650611 | Pseudomonadaceae | Pseudomonas_E_647464 | |
| Sun start vs. Shade start | Shade start | 4.017 | -19.623 | 0.004 | 0.0466 | ASV200 | Bacteria | Pseudomonadota | Alphaproteobacteria | Caulobacterales | TH1-2 | Vitreimonas | Vitreimonas sp001464425 |
| Sun start vs. Shade start | Shade start | 263.592 | -19.56 | 0.0041 | 0.0468 | ASV374 | Bacteria | Bacillota_I | Bacilli_A | Bacillales_B_310392 | Bacillaceae_G | Bacillus_A | |
| Sun UV vs. Sun noUV | Sun noUV | 19.2 | -48.341 | <0.0001 | <0.0001 | ASV97 | Bacteria | Pseudomonadota | Gammaproteobacteria | Enterobacterales_737866 | Enterobacteriaceae_A_726016 | Gibbsiella | Gibbsiella greigii |
| Sun UV vs. Sun noUV | Sun UV | 2.695 | 47.112 | <0.0001 | <0.0001 | ASV379 | Bacteria | Acidobacteriota | Terriglobia | Bryobacterales | Bryobacteraceae | KBS-96 | KBS-96 sp000381625 |
| Sun UV vs. Sun noUV | Sun noUV | 35.122 | -42.323 | <0.0001 | <0.0001 | ASV47 | Bacteria | Pseudomonadota | Alphaproteobacteria | Rhizobiales_505101 | Beijerinckiaceae | Methylobacterium | Methylobacterium sp001424705 |
| Sun UV vs. Sun noUV | Sun UV | 30.609 | 44.221 | <0.0001 | <0.0001 | ASV115 | Bacteria | Actinomycetota | Actinomycetes | Actinomycetales | Micrococcaceae | Micrococcus | Micrococcus luteus |

| | | | | | | | | | | | | | |
|---|---|---|---|---|---|---|---|---|---|---|---|---|---|
| Sun UV vs. Sun noUV | Sun UV | 15.816 | 42.141 | <0.0001 | <0.0001 | ASV397 | Bacteria | Bacillota_I | Bacilli_A | Lactobacillales | Streptococcaceae | Streptococcus | |
| Sun UV vs. Sun noUV | Sun noUV | 4.481 | -41.613 | <0.0001 | <0.0001 | ASV93 | Bacteria | Pseudomonadota | Alphaproteobacteria | Rhizobiales_505101 | Rhizobiaceae | | |
| Sun UV vs. Sun noUV | Sun UV | 2.965 | 41.045 | <0.0001 | <0.0001 | ASV341 | Bacteria | Pseudomonadota | Alphaproteobacteria | | | | |
| Sun UV vs. Sun noUV | Sun UV | 78.456 | 25.111 | <0.0001 | <0.0001 | ASV61 | Bacteria | Desulfobacterota_G_459546 | Desulfuromonadia | Geobacterales | Geobacteraceae_439684 | | |
| Sun UV vs. Sun noUV | Sun UV | 263.592 | 39.242 | <0.0001 | <0.0001 | ASV374 | Bacteria | Bacillota_I | Bacilli_A | Bacillales_B_310392 | Bacillaceae_G | Bacillus_A | |
| Sun UV vs. Sun noUV | Sun UV | 2.437 | 37.773 | <0.0001 | <0.0001 | ASV56 | Bacteria | Bacteroidota | Bacteroidia | Cytophagales_B | Hymenobacteraceae | Solirubrum | |
| Sun UV vs. Sun noUV | Sun UV | 91.323 | 37.266 | <0.0001 | <0.0001 | ASV351 | Bacteria | Bacillota_I | Bacilli_A | Bacillales_B_306087 | DSM-18226_301387 | | |
| Sun UV vs. Sun noUV | Sun UV | 18.308 | 36.653 | <0.0001 | <0.0001 | ASV119 | Bacteria | Pseudomonadota | Alphaproteobacteria | Acetobacterales | Acetobacteraceae | Roseomonas_507234 | Roseomonas sp001941945 |
| Sun UV vs. Sun noUV | Sun noUV | 103.969 | -27.114 | <0.0001 | <0.0001 | ASV30 | Bacteria | Pseudomonadota | Gammaproteobacteria | Enterobacterales_737866 | Enterobacteriaceae_A_729055 | Yersinia | Lonsdalea iberica |
| Sun UV vs. Sun noUV | Sun noUV | 33.421 | -25.732 | <0.0001 | <0.0001 | ASV43 | Bacteria | Pseudomonadota | Alphaproteobacteria | Sphingomonadales | Sphingomonadaceae_486827 | Sphingomonas_L_486704 | |
| Sun UV vs. Sun noUV | Sun noUV | 48.705 | -24.993 | <0.0001 | <0.0001 | ASV62 | Bacteria | Pseudomonadota | Alphaproteobacteria | Rhizobiales_505101 | Beijerinckiaceae | Lichenihabitans_504419 | Lichenihabitans ramalinae |
| Sun UV vs. Sun noUV | Sun noUV | 3.012 | -34.16 | <0.0001 | <0.0001 | ASV88 | Bacteria | Actinomycetota | Actinomycetes | Mycobacteriales | Mycobacteriaceae | Lawsonella | |
| Sun UV vs. Sun noUV | Sun UV | 3.112 | 33.716 | <0.0001 | <0.0001 | ASV206 | Bacteria | Actinomycetota | Actinomycetes | Actinomycetales | Dermatophilaceae | | |
| Sun UV vs. Sun noUV | Sun noUV | 20.003 | -33.492 | <0.0001 | <0.0001 | ASV172 | Bacteria | Bacillota_I | Bacilli_A | Lactobacillales | Streptococcaceae | Lactococcus_A_346120 | |
| Sun UV vs. Sun noUV | Sun UV | 19.437 | 31.882 | <0.0001 | <0.0001 | ASV305 | Bacteria | Pseudomonadota | Alphaproteobacteria | Rhodobacterales | Rhodobacteraceae | | |
| Sun UV vs. Sun noUV | Sun UV | 9.519 | 31.67 | <0.0001 | <0.0001 | ASV214 | Bacteria | Bacteroidota | Bacteroidia | Flavobacteriales_B_877923 | Weeksellaceae | | |
| Sun UV vs. Sun noUV | Sun UV | 4.051 | 31.632 | <0.0001 | <0.0001 | ASV495 | Bacteria | Bacillota_I | Bacilli_A | Paenibacillales | Paenibacillaceae_367444 | Saccharibacillus | Saccharibacillus sacchari |
| Sun UV vs. Sun noUV | Sun UV | 23.836 | 30.501 | <0.0001 | <0.0001 | ASV634 | Bacteria | Bacillota_A_368345 | Clostridia_258483 | Oscillospirales | Ethanoligenenaceae | Ruminiclostridium_D | Ruminiclostridium_D cellulosi |
| Sun UV vs. Sun noUV | Sun UV | 2.39 | 30.266 | <0.0001 | <0.0001 | ASV795 | Bacteria | Pseudomonadota | Alphaproteobacteria | Acetobacterales | Acetobacteraceae | | |
| Sun UV vs. Sun noUV | Sun UV | 2.064 | 28.693 | <0.0001 | <0.0001 | ASV329 | Bacteria | Bacteroidota | Bacteroidia | Flavobacteriales_B_877923 | Weeksellaceae | | |
| Sun UV vs. Sun noUV | Sun noUV | 10.132 | -28.426 | <0.0001 | <0.0001 | ASV193 | Bacteria | Bacteroidota | Bacteroidia | Sphingobacteriales | Sphingobacteriaceae | Pedobacter_B_887417 | |
| Sun UV vs. Sun noUV | Sun UV | 20.423 | 28.032 | <0.0001 | <0.0001 | ASV252 | Bacteria | Pseudomonadota | Alphaproteobacteria | Rhodobacterales | Rhodobacteraceae | Paracoccus | |
| Sun UV vs. Sun noUV | Sun UV | 8.234 | 27.852 | <0.0001 | <0.0001 | ASV948 | Bacteria | Pseudomonadota | Alphaproteobacteria | Rhizobiales_505101 | Beijerinckiaceae | Methylobacterium | Methylobacterium radiotolerans |

| | | | | | | | | | | | | | |
|---|---|---|---|---|---|---|---|---|---|---|---|---|---|
| Sun UV vs. Sun noUV | Sun noUV | 4.889 | -27.9 | <0.0001 | <0.0001 | ASV180 | Bacteria | Pseudomonadota | Alphaproteobacteria | Sphingomonadales | Sphingomonadaceae_486827 | Sphingomonas_L_486704 | Sphingomonas_L_486704 aracearum |
| Sun UV vs. Sun noUV | Sun UV | 10.421 | 27.784 | <0.0001 | <0.0001 | ASV1384 | Bacteria | Pseudomonadota | Alphaproteobacteria | Rhizobiales_505101 | Beijerinckiaceae | Methylobacterium | |
| Sun UV vs. Sun noUV | Sun UV | 6.586 | 27.743 | <0.0001 | <0.0001 | ASV155 | Bacteria | Pseudomonadota | Gammaproteobacteria | Burkholderiales | Burkholderiaceae_A_595421 | Cupriavidus | |
| Sun UV vs. Sun noUV | Sun UV | 8.524 | 27.624 | <0.0001 | <0.0001 | ASV169 | Bacteria | Acidobacteriota | Terriglobia | Terriglobales | Acidobacteriaceae | | |
| Sun UV vs. Sun noUV | Sun UV | 6.394 | 27.379 | <0.0001 | <0.0001 | ASV564 | Bacteria | Pseudomonadota | Gammaproteobacteria | Pseudomonadales_A_637066 | Oleiphilaceae_637066 | Halospina_636536 | Halospina utahensis_B |
| Sun UV vs. Sun noUV | Sun noUV | 10.012 | -27.06 | <0.0001 | <0.0001 | ASV84 | Bacteria | Pseudomonadota | Alphaproteobacteria | Sphingomonadales | Sphingomonadaceae_486827 | Sphingomonas_L_486704 | |
| Sun UV vs. Sun noUV | Sun UV | 3.615 | 26.972 | <0.0001 | <0.0001 | ASV655 | Bacteria | Actinomycetota | Thermoleophilia | Solirubrobacterales | Solirubrobacteraceae | Solirubrobacter | Solirubrobacter soli |
| Sun UV vs. Sun noUV | Sun UV | 16.893 | 26.813 | <0.0001 | <0.0001 | ASV345 | Bacteria | Bacillota_I | Bacilli_A | Paenibacillales | Paenibacillaceae_367444 | Saccharibacillus | Saccharibacillus sacchari |
| Sun UV vs. Sun noUV | Sun noUV | 7.867 | -26.795 | <0.0001 | <0.0001 | ASV194 | Bacteria | Actinomycetota | Actinomycetes | Streptosporangiales | Streptosporangiaceae | Actinoallomurus | |
| Sun UV vs. Sun noUV | Sun noUV | 4.741 | -26.608 | <0.0001 | <0.0001 | ASV457 | Bacteria | Pseudomonadota | Alphaproteobacteria | Acetobacterales | Acetobacteraceae | | |
| Sun UV vs. Sun noUV | Sun UV | 14.311 | 22.082 | <0.0001 | <0.0001 | ASV105 | Bacteria | Actinomycetota | Actinomycetes | Actinomycetales | Microbacteriaceae | | |
| Sun UV vs. Sun noUV | Sun UV | 10.655 | 25.965 | <0.0001 | <0.0001 | ASV520 | Bacteria | Actinomycetota | Actinomycetes | Actinomycetales | Brevibacteriaceae | Brevibacterium | |
| Sun UV vs. Sun noUV | Sun UV | 2.768 | 25.969 | <0.0001 | <0.0001 | ASV1075 | Bacteria | Bacteroidota | Bacteroidia | Flavobacteriales_B_877923 | Weeksellaceae | Kaistella | Kaistella sp000023725 |
| Sun UV vs. Sun noUV | Sun noUV | 16.539 | -23.824 | <0.0001 | <0.0001 | ASV137 | Bacteria | Actinomycetota | Actinomycetes | Mycobacteriales | Mycobacteriaceae | Rhodococcus_376336 | |
| Sun UV vs. Sun noUV | Sun UV | 22.221 | 25.025 | <0.0001 | <0.0001 | ASV154 | Bacteria | Actinomycetota | Actinomycetes | Actinomycetales | | | |
| Sun UV vs. Sun noUV | Sun noUV | 30.844 | -24.975 | <0.0001 | <0.0001 | ASV118 | Bacteria | Actinomycetota | Actinomycetes | Mycobacteriales | Mycobacteriaceae | Corynebacterium | |
| Sun UV vs. Sun noUV | Sun noUV | 11.174 | -24.04 | <0.0001 | <0.0001 | ASV66 | Bacteria | Pseudomonadota | Alphaproteobacteria | Sphingomonadales | Sphingomonadaceae_486827 | Sphingomonas_N_483429 | |
| Sun UV vs. Sun noUV | Sun UV | 113.399 | 19.77 | <0.0001 | 0.0001 | ASV332 | Bacteria | Pseudomonadota | Gammaproteobacteria | Burkholderiales | Burkholderiaceae_A_595421 | | |
| Shade UV vs. Shade noUV | Shade UV | 13.277 | 45.764 | <0.0001 | <0.0001 | ASV134 | Bacteria | Actinomycetota | Actinomycetes | Mycobacteriales | Micromonosporaceae | | |
| Shade UV vs. Shade noUV | Shade noUV | 8.768 | -47.764 | <0.0001 | <0.0001 | ASV75 | Bacteria | Bacteroidota | Bacteroidia | Cytophagales_B | Hymenobacteraceae | Hymenobacter_910554 | Hymenobacter setariae |
| Shade UV vs. Shade noUV | Shade noUV | 4.017 | -46.473 | <0.0001 | <0.0001 | ASV200 | Bacteria | Pseudomonadota | Alphaproteobacteria | Caulobacterales | TH1-2 | Vitreimonas | Vitreimonas sp001464425 |
| Shade UV vs. Shade noUV | Shade noUV | 15.816 | -44.028 | <0.0001 | <0.0001 | ASV397 | Bacteria | Bacillota_I | Bacilli_A | Lactobacillales | Streptococcaceae | Streptococcus | |
| Shade UV vs. Shade noUV | Shade UV | 4.481 | 42.715 | <0.0001 | <0.0001 | ASV93 | Bacteria | Pseudomonadota | Alphaproteobacteria | Rhizobiales_505101 | Rhizobiaceae | | |

| | | | | | | | | | | | | | |
|---|---|---|---|---|---|---|---|---|---|---|---|---|---|
| Shade UV vs. Shade noUV | Shade UV | 18.308 | 42.122 | <0.0001 | <0.0001 | ASV119 | Bacteria | Pseudomonadota | Alphaproteobacteria | Acetobacterales | Acetobacteraceae | Roseomonas_507234 | Roseomonas sp001941945 |
| Shade UV vs. Shade noUV | Shade UV | 2.965 | 41.633 | <0.0001 | <0.0001 | ASV341 | Bacteria | Pseudomonadota | Alphaproteobacteria | | | | |
| Shade UV vs. Shade noUV | Shade noUV | 371.071 | -28.905 | <0.0001 | <0.0001 | ASV96 | Bacteria | Pseudomonadota | Gammaproteobacteria | Pseudomonadales_A_650611 | Pseudomonadaceae | Pseudomonas_E_647464 | |
| Shade UV vs. Shade noUV | Shade noUV | 20.003 | -41.211 | <0.0001 | <0.0001 | ASV172 | Bacteria | Bacillota_I | Bacilli_A | Lactobacillales | Streptococcaceae | Lactococcus_A_346120 | |
| Shade UV vs. Shade noUV | Shade UV | 29.691 | 38.669 | <0.0001 | <0.0001 | ASV270 | Bacteria | Pseudomonadota | Gammaproteobacteria | Pseudomonadales_A_650611 | Pseudomonadaceae | Pseudomonas_E_647464 | |
| Shade UV vs. Shade noUV | Shade UV | 2.437 | 37.537 | <0.0001 | <0.0001 | ASV56 | Bacteria | Bacteroidota | Bacteroidia | Cytophagales_B | Hymenobacteraceae | Solirubrum | |
| Shade UV vs. Shade noUV | Shade UV | 32.122 | 35.941 | <0.0001 | <0.0001 | ASV213 | Bacteria | Actinomycetota | Actinomycetes | Propionibacteriales | Nocardioidaceae | Nocardioides_A_392796 | |
| Shade UV vs. Shade noUV | Shade UV | 13.048 | 26.829 | <0.0001 | <0.0001 | ASV107 | Bacteria | Pseudomonadota | Alphaproteobacteria | Acetobacterales | Acetobacteraceae | | |
| Shade UV vs. Shade noUV | Shade UV | 33.421 | 23.779 | <0.0001 | <0.0001 | ASV43 | Bacteria | Pseudomonadota | Alphaproteobacteria | Sphingomonadales | Sphingomonadaceae_486827 | Sphingomonas_L_486704 | |
| Shade UV vs. Shade noUV | Shade UV | 5.876 | 32.301 | <0.0001 | <0.0001 | ASV277 | Bacteria | Pseudomonadota | Gammaproteobacteria | Burkholderiales | Burkholderiaceae_A_595421 | | |
| Shade UV vs. Shade noUV | Shade UV | 16.537 | 27.35 | <0.0001 | <0.0001 | ASV114 | Bacteria | Pseudomonadota | Alphaproteobacteria | Rhizobiales_505101 | | | |
| Shade UV vs. Shade noUV | Shade noUV | 6.394 | -31.14 | <0.0001 | <0.0001 | ASV564 | Bacteria | Pseudomonadota | Gammaproteobacteria | Pseudomonadales_A_637066 | Oleiphilaceae_637066 | Halospina_636536 | Halospina utahensis_B |
| Shade UV vs. Shade noUV | Shade UV | 46.867 | 24.021 | <0.0001 | <0.0001 | ASV65 | Bacteria | Bacteroidota | Bacteroidia | Sphingobacteriales | Sphingobacteriaceae | Pedobacter_B_887417 | |
| Shade UV vs. Shade noUV | Shade UV | 2.896 | 28.745 | <0.0001 | <0.0001 | ASV199 | Bacteria | Actinomycetota | Actinomycetes | Propionibacteriales | Nocardioidaceae | Nocardioides_A_392796 | |
| Shade UV vs. Shade noUV | Shade UV | 4.62 | 28.46 | <0.0001 | <0.0001 | ASV163 | Bacteria | Pseudomonadota | Gammaproteobacteria | Burkholderiales | Burkholderiaceae_A_595421 | Caballeronia | Caballeronia udeis |
| Shade UV vs. Shade noUV | Shade UV | 3.615 | 28.358 | <0.0001 | <0.0001 | ASV655 | Bacteria | Actinomycetota | Thermoleophilia | Solirubrobacterales | Solirubrobacteraceae | Solirubrobacter | Solirubrobacter soli |
| Shade UV vs. Shade noUV | Shade UV | 13.145 | 28.021 | <0.0001 | <0.0001 | ASV179 | Bacteria | Actinomycetota | Actinomycetes | Streptomycetales_400645 | Streptomycetaceae_400641 | Streptomyces_400150 | |
| Shade UV vs. Shade noUV | Shade UV | 8.524 | 27.545 | <0.0001 | <0.0001 | ASV169 | Bacteria | Acidobacteriota | Terriglobia | Terriglobales | Acidobacteriaceae | | |
| Shade UV vs. Shade noUV | Shade UV | 14.311 | 22.989 | <0.0001 | <0.0001 | ASV105 | Bacteria | Actinomycetota | Actinomycetes | Actinomycetales | Microbacteriaceae | | |
| Shade UV vs. Shade noUV | Shade UV | 24.338 | 26.964 | <0.0001 | <0.0001 | ASV222 | Bacteria | Pseudomonadota | Gammaproteobacteria | Burkholderiales | Burkholderiaceae_A_595421 | | |
| Shade UV vs. Shade noUV | Shade noUV | 1.827 | -26.398 | <0.0001 | <0.0001 | ASV335 | Bacteria | Actinomycetota | Actinomycetes | Actinomycetales | Brevibacteriaceae | Brevibacterium | |
| Shade UV vs. Shade noUV | Shade UV | 17.859 | 25.958 | <0.0001 | <0.0001 | ASV92 | Bacteria | Pseudomonadota | Gammaproteobacteria | Burkholderiales | Burkholderiaceae_A_595421 | | |
| Shade UV vs. Shade noUV | Shade UV | 5.173 | 25.81 | <0.0001 | <0.0001 | ASV130 | Bacteria | Pseudomonadota | Alphaproteobacteria | Sphingomonadales | Sphingomonadaceae_486827 | | |

| | | | | | | | | | | | | | |
|---|---|---|---|---|---|---|---|---|---|---|---|---|---|
| Shade UV vs. Shade noUV | Shade noUV | 10.012 | -25.432 | <0.0001 | <0.0001 | ASV84 | Bacteria | Pseudomonadota | Alphaproteobacteria | Sphingomonadales | Sphingomonadaceae_486827 | Sphingomonas_L_486704 | |
| Shade UV vs. Shade noUV | Shade noUV | 4.889 | -25.265 | <0.0001 | <0.0001 | ASV180 | Bacteria | Pseudomonadota | Alphaproteobacteria | Sphingomonadales | Sphingomonadaceae_486827 | Sphingomonas_L_486704 | Sphingomonas_L_486704 aracearum |
| Shade UV vs. Shade noUV | Shade UV | 4.191 | 25.105 | <0.0001 | <0.0001 | ASV2012 | Bacteria | Pseudomonadota | Alphaproteobacteria | Sphingomonadales | Sphingomonadaceae_486827 | Sphingomonas_L_486704 | Sphingomonas_L_486704 mali |
| Shade UV vs. Shade noUV | Shade UV | 9.238 | 21.746 | <0.0001 | <0.0001 | ASV86 | Bacteria | Pseudomonadota | Alphaproteobacteria | Caulobacterales | Caulobacteraceae | Caulobacter_487784 | |
| Shade UV vs. Shade noUV | Shade UV | 22.44 | 23.941 | <0.0001 | <0.0001 | ASV216 | Bacteria | Bacillota_I | Bacilli_A | Paenibacillales | Paenibacillaceae_367444 | | |
| Shade UV vs. Shade noUV | Shade noUV | 5.253 | -24.882 | <0.0001 | <0.0001 | ASV159 | Bacteria | Pseudomonadota | Alphaproteobacteria | Rhizobiales_505101 | Beijerinckiaceae | 4M-Z18 | 4M-Z18 sp003400305 |
| Shade UV vs. Shade noUV | Shade noUV | 14.542 | -23.796 | <0.0001 | <0.0001 | ASV294 | Bacteria | Pseudomonadota | Alphaproteobacteria | Rhizobiales_505101 | Beijerinckiaceae | Microvirga | |
| Shade UV vs. Shade noUV | Shade UV | 5.132 | 23.65 | <0.0001 | <0.0001 | ASV203 | Bacteria | Pseudomonadota | Alphaproteobacteria | Acetobacterales | Acetobacteraceae | | |
| Shade UV vs. Shade noUV | Shade UV | 10.655 | 23.493 | <0.0001 | <0.0001 | ASV520 | Bacteria | Actinomycetota | Actinomycetes | Actinomycetales | Brevibacteriaceae | Brevibacterium | |
| Shade UV vs. Shade noUV | Shade UV | 4.294 | 22.322 | <0.0001 | <0.0001 | ASV413 | Bacteria | Actinomycetota | Thermoleophilia | Solirubrobacterales | Solirubrobacteraceae | Baekduia | Baekduia soli |
| Shade UV vs. Shade noUV | Shade UV | 3.952 | 22.26 | <0.0001 | <0.0001 | ASV224 | Bacteria | Pseudomonadota | Alphaproteobacteria | Sphingomonadales | Sphingomonadaceae_486827 | | |
| Shade UV vs. Shade noUV | Shade UV | 12.569 | 21.603 | <0.0001 | 0.0002 | ASV703 | Bacteria | Pseudomonadota | Alphaproteobacteria | Rhizobiales_505101 | Beijerinckiaceae | | |
| Shade UV vs. Shade noUV | Shade noUV | 213.321 | -11.805 | <0.0001 | 0.0003 | ASV20 | Bacteria | Pseudomonadota | Alphaproteobacteria | Rhizobiales_505101 | Beijerinckiaceae | Methylobacterium | Methylobacterium brachiatum |
| Shade UV vs. Shade noUV | Shade UV | 809.259 | 8.066 | 0.0003 | 0.0037 | ASV10 | Bacteria | Pseudomonadota | Gammaproteobacteria | Pseudomonadales_A_650611 | Pseudomonadaceae | Pseudomonas_E_647464 | |
| Sun UV vs. Shade noUV | Shade noUV | 19.2 | -50.491 | <0.0001 | <0.0001 | ASV97 | Bacteria | Pseudomonadota | Gammaproteobacteria | Enterobacterales_737866 | Enterobacteriaceae_A_726016 | Gibbsiella | Gibbsiella greigii |
| Sun UV vs. Shade noUV | Sun UV | 13.277 | 44.111 | <0.0001 | <0.0001 | ASV134 | Bacteria | Actinomycetota | Actinomycetes | Mycobacteriales | Micromonosporaceae | | |
| Sun UV vs. Shade noUV | Shade noUV | 35.122 | -39.931 | <0.0001 | <0.0001 | ASV47 | Bacteria | Pseudomonadota | Alphaproteobacteria | Rhizobiales_505101 | Beijerinckiaceae | Methylobacterium | Methylobacterium sp001424705 |
| Sun UV vs. Shade noUV | Sun UV | 2.965 | 41.521 | <0.0001 | <0.0001 | ASV341 | Bacteria | Pseudomonadota | Alphaproteobacteria | | | | |
| Sun UV vs. Shade noUV | Shade noUV | 20.003 | -40.092 | <0.0001 | <0.0001 | ASV172 | Bacteria | Bacillota_I | Bacilli_A | Lactobacillales | Streptococcaceae | Lactococcus_A_346120 | |
| Sun UV vs. Shade noUV | Shade noUV | 103.969 | -29.564 | <0.0001 | <0.0001 | ASV30 | Bacteria | Pseudomonadota | Gammaproteobacteria | Enterobacterales_737866 | Enterobacteriaceae_A_729055 | Yersinia | Lonsdalea iberica |
| Sun UV vs. Shade noUV | Sun UV | 263.592 | 39.614 | <0.0001 | <0.0001 | ASV374 | Bacteria | Bacillota_I | Bacilli_A | Bacillales_B_310392 | Bacillaceae_G | Bacillus_A | |
| Sun UV vs. Shade noUV | Sun UV | 2.437 | 38.341 | <0.0001 | <0.0001 | ASV56 | Bacteria | Bacteroidota | Bacteroidia | Cytophagales_B | Hymenobacteraceae | Solirubrum | |
| Sun UV vs. Shade noUV | Shade noUV | 3.012 | -37.64 | <0.0001 | <0.0001 | ASV88 | Bacteria | Actinomycetota | Actinomycetes | Mycobacteriales | Mycobacteriaceae | Lawsonella | |

| | | | | | | | | | | | | | |
|---|---|---|---|---|---|---|---|---|---|---|---|---|---|
| Sun UV vs. Shade noUV | Sun UV | 91.323 | 37.592 | <0.0001 | <0.0001 | ASV351 | Bacteria | Bacillota_I | Bacilli_A | Bacillales_B_306087 | DSM-18226_301387 | | |
| Sun UV vs. Shade noUV | Shade noUV | 14.928 | -27.187 | <0.0001 | <0.0001 | ASV117 | Bacteria | Pseudomonadota | Alphaproteobacteria | Caulobacterales | Caulobacteraceae | | |
| Sun UV vs. Shade noUV | Sun UV | 13.048 | 27.774 | <0.0001 | <0.0001 | ASV107 | Bacteria | Pseudomonadota | Alphaproteobacteria | Acetobacterales | Acetobacteraceae | | |
| Sun UV vs. Shade noUV | Sun UV | 18.308 | 36.954 | <0.0001 | <0.0001 | ASV119 | Bacteria | Pseudomonadota | Alphaproteobacteria | Acetobacterales | Acetobacteraceae | Roseomonas_507234 | Roseomonas sp001941945 |
| Sun UV vs. Shade noUV | Sun UV | 66.071 | 35.931 | <0.0001 | <0.0001 | ASV150 | Bacteria | Actinomycetota | Actinomycetes | Propionibacteriales | Nocardioidaceae | Nocardioides_A_392796 | |
| Sun UV vs. Shade noUV | Sun UV | 40.002 | 35.665 | <0.0001 | <0.0001 | ASV34 | Bacteria | Pseudomonadota | Gammaproteobacteria | Enterobacterales_737866 | Enterobacteriaceae_A_725029 | Pluralibacter_724998 | |
| Sun UV vs. Shade noUV | Sun UV | 3.112 | 34.086 | <0.0001 | <0.0001 | ASV206 | Bacteria | Actinomycetota | Actinomycetes | Actinomycetales | Dermatophilaceae | | |
| Sun UV vs. Shade noUV | Sun UV | 9.519 | 32.301 | <0.0001 | <0.0001 | ASV214 | Bacteria | Bacteroidota | Bacteroidia | Flavobacteriales_B_877923 | Weeksellaceae | | |
| Sun UV vs. Shade noUV | Sun UV | 4.051 | 31.883 | <0.0001 | <0.0001 | ASV495 | Bacteria | Bacillota_I | Bacilli_A | Paenibacillales | Paenibacillaceae_367444 | Saccharibacillus | Saccharibacillus sacchari |
| Sun UV vs. Shade noUV | Sun UV | 46.867 | 24.512 | <0.0001 | <0.0001 | ASV65 | Bacteria | Bacteroidota | Bacteroidia | Sphingobacteriales | Sphingobacteriaceae | Pedobacter_B_887417 | |
| Sun UV vs. Shade noUV | Sun UV | 19.437 | 31.529 | <0.0001 | <0.0001 | ASV305 | Bacteria | Pseudomonadota | Alphaproteobacteria | Rhodobacterales | Rhodobacteraceae | | |
| Sun UV vs. Shade noUV | Shade noUV | 48.705 | -22.125 | <0.0001 | <0.0001 | ASV62 | Bacteria | Pseudomonadota | Alphaproteobacteria | Rhizobiales_505101 | Beijerinckiaceae | Lichenihabitans_504419 | Lichenihabitans ramalinae |
| Sun UV vs. Shade noUV | Sun UV | 23.836 | 30.825 | <0.0001 | <0.0001 | ASV634 | Bacteria | Bacillota_A_368345 | Clostridia_258483 | Oscillospirales | Ethanoligenenaceae | Ruminiclostridium_D | Ruminiclostridium_D cellulosi |
| Sun UV vs. Shade noUV | Sun UV | 2.39 | 30.703 | <0.0001 | <0.0001 | ASV795 | Bacteria | Pseudomonadota | Alphaproteobacteria | Acetobacterales | Acetobacteraceae | | |
| Sun UV vs. Shade noUV | Shade noUV | 16.539 | -27.777 | <0.0001 | <0.0001 | ASV137 | Bacteria | Actinomycetota | Actinomycetes | Mycobacteriales | Mycobacteriaceae | Rhodococcus_376336 | |
| Sun UV vs. Shade noUV | Sun UV | 16.537 | 25.952 | <0.0001 | <0.0001 | ASV114 | Bacteria | Pseudomonadota | Alphaproteobacteria | Rhizobiales_505101 | | | |
| Sun UV vs. Shade noUV | Sun UV | 2.064 | 29.365 | <0.0001 | <0.0001 | ASV329 | Bacteria | Bacteroidota | Bacteroidia | Flavobacteriales_B_877923 | Weeksellaceae | | |
| Sun UV vs. Shade noUV | Sun UV | 20.423 | 28.583 | <0.0001 | <0.0001 | ASV252 | Bacteria | Pseudomonadota | Alphaproteobacteria | Rhodobacterales | Rhodobacteraceae | Paracoccus | |
| Sun UV vs. Shade noUV | Sun UV | 13.145 | 28.039 | <0.0001 | <0.0001 | ASV179 | Bacteria | Actinomycetota | Actinomycetes | Streptomycetales_400645 | Streptomycetaceae_400641 | Streptomyces_400150 | |
| Sun UV vs. Shade noUV | Sun UV | 8.524 | 27.891 | <0.0001 | <0.0001 | ASV169 | Bacteria | Acidobacteriota | Terriglobia | Terriglobales | Acidobacteriaceae | | |
| Sun UV vs. Shade noUV | Sun UV | 8.234 | 27.886 | <0.0001 | <0.0001 | ASV948 | Bacteria | Pseudomonadota | Alphaproteobacteria | Rhizobiales_505101 | Beijerinckiaceae | Methylobacterium | Methylobacterium radiotolerans |
| Sun UV vs. Shade noUV | Sun UV | 10.421 | 27.886 | <0.0001 | <0.0001 | ASV1384 | Bacteria | Pseudomonadota | Alphaproteobacteria | Rhizobiales_505101 | Beijerinckiaceae | Methylobacterium | |
| Sun UV vs. Shade noUV | Sun UV | 6.586 | 27.608 | <0.0001 | <0.0001 | ASV155 | Bacteria | Pseudomonadota | Gammaproteobacteria | Burkholderiales | Burkholderiaceae_A_595421 | Cupriavidus | |

| | | | | | | | | | | | | | |
|---|---|---|---|---|---|---|---|---|---|---|---|---|---|
| Sun UV vs. Shade noUV | Sun UV | 3.615 | 27.41 | <0.0001 | <0.0001 | ASV655 | Bacteria | Actinomycetota | Thermoleophilia | Solirubrobacterales | Solirubrobacteraceae | Solirubrobacter | Solirubrobacter soli |
| Sun UV vs. Shade noUV | Shade noUV | 4.889 | -27.249 | <0.0001 | <0.0001 | ASV180 | Bacteria | Pseudomonadota | Alphaproteobacteria | Sphingomonadales | Sphingomonadaceae_486827 | Sphingomonas_L_486704 | Sphingomonas_L_486704 aracearum |
| Sun UV vs. Shade noUV | Sun UV | 14.311 | 22.523 | <0.0001 | <0.0001 | ASV105 | Bacteria | Actinomycetota | Actinomycetes | Actinomycetales | Microbacteriaceae | | |
| Sun UV vs. Shade noUV | Shade noUV | 1.827 | -26.599 | <0.0001 | <0.0001 | ASV335 | Bacteria | Actinomycetota | Actinomycetes | Actinomycetales | Brevibacteriaceae | Brevibacterium | |
| Sun UV vs. Shade noUV | Sun UV | 10.655 | 26.594 | <0.0001 | <0.0001 | ASV520 | Bacteria | Actinomycetota | Actinomycetes | Actinomycetales | Brevibacteriaceae | Brevibacterium | |
| Sun UV vs. Shade noUV | Shade noUV | 10.012 | -26.5 | <0.0001 | <0.0001 | ASV84 | Bacteria | Pseudomonadota | Alphaproteobacteria | Sphingomonadales | Sphingomonadaceae_486827 | Sphingomonas_L_486704 | |
| Sun UV vs. Shade noUV | Shade noUV | 5.253 | -26.557 | <0.0001 | <0.0001 | ASV159 | Bacteria | Pseudomonadota | Alphaproteobacteria | Rhizobiales_505101 | Beijerinckiaceae | 4M-Z18 | 4M-Z18 sp003400305 |
| Sun UV vs. Shade noUV | Shade noUV | 5.889 | -26.558 | <0.0001 | <0.0001 | ASV803 | Bacteria | Pseudomonadota | Gammaproteobacteria | Burkholderiales | Neisseriaceae | Snodgrassella | Snodgrassella alvi |
| Sun UV vs. Shade noUV | Shade noUV | 58.059 | -26.383 | <0.0001 | <0.0001 | ASV102 | Bacteria | Actinomycetota | Actinomycetes | Mycobacteriales | Pseudonocardiaceae | Actinomycetospora | |
| Sun UV vs. Shade noUV | Sun UV | 4.62 | 26.263 | <0.0001 | <0.0001 | ASV163 | Bacteria | Pseudomonadota | Gammaproteobacteria | Burkholderiales | Burkholderiaceae_A_595421 | Caballeronia | Caballeronia udeis |
| Sun UV vs. Shade noUV | Sun UV | 2.768 | 26.237 | <0.0001 | <0.0001 | ASV1075 | Bacteria | Bacteroidota | Bacteroidia | Flavobacteriales_B_877923 | Weeksellaceae | Kaistella | Kaistella sp000023725 |
| Sun UV vs. Shade noUV | Shade noUV | 10.132 | -26.129 | <0.0001 | <0.0001 | ASV193 | Bacteria | Bacteroidota | Bacteroidia | Sphingobacteriales | Sphingobacteriaceae | Pedobacter_B_887417 | |
| Sun UV vs. Shade noUV | Sun UV | 7.23 | 25.969 | <0.0001 | <0.0001 | ASV404 | Bacteria | Bacteroidota | Bacteroidia | Sphingobacteriales | Sphingobacteriaceae | Mucilaginibacter_A | |
| Sun UV vs. Shade noUV | Sun UV | 9.11 | 23.334 | <0.0001 | <0.0001 | ASV146 | Bacteria | Pseudomonadota | Alphaproteobacteria | Acetobacterales | Acetobacteraceae | | |
| Sun UV vs. Shade noUV | Sun UV | 9.238 | 21.729 | <0.0001 | <0.0001 | ASV86 | Bacteria | Pseudomonadota | Alphaproteobacteria | Caulobacterales | Caulobacteraceae | Caulobacter_487784 | |
| Sun UV vs. Shade noUV | Sun UV | 22.221 | 24.665 | <0.0001 | <0.0001 | ASV154 | Bacteria | Actinomycetota | Actinomycetes | Actinomycetales | | | |
| Sun UV vs. Shade noUV | Shade noUV | 14.542 | -24.344 | <0.0001 | <0.0001 | ASV294 | Bacteria | Pseudomonadota | Alphaproteobacteria | Rhizobiales_505101 | Beijerinckiaceae | Microvirga | |
| Sun UV vs. Shade noUV | Sun UV | 22.44 | 22.718 | <0.0001 | <0.0001 | ASV216 | Bacteria | Bacillota_I | Bacilli_A | Paenibacillales | Paenibacillaceae_367444 | | |
| Sun UV vs. Shade noUV | Sun UV | 5.097 | 23.309 | <0.0001 | <0.0001 | ASV307 | Bacteria | Pseudomonadota | Gammaproteobacteria | Enterobacterales_737866 | Enterobacteriaceae_A_680892 | Izhakiella | Duffyella gerundensis |
| Sun UV vs. Shade noUV | Shade noUV | 30.844 | -22.297 | <0.0001 | <0.0001 | ASV118 | Bacteria | Actinomycetota | Actinomycetes | Mycobacteriales | Mycobacteriaceae | Corynebacterium | |
| Sun UV vs. Shade noUV | Shade noUV | 213.321 | -12.052 | <0.0001 | <0.0001 | ASV20 | Bacteria | Pseudomonadota | Alphaproteobacteria | Rhizobiales_505101 | Beijerinckiaceae | Methylobacterium | Methylobacterium brachiatum |
| Control UV vs. Control noUV | Control UV | 431.076 | 29.011 | <0.0001 | <0.0001 | ASV29 | Bacteria | Actinomycetota | Actinomycetes | Actinomycetales | Microbacteriaceae | | |
| Control UV vs. Control noUV | Control UV | 100.15 | 21.257 | <0.0001 | 0.0002 | ASV67 | Bacteria | Pseudomonadota | Gammaproteobacteria | Burkholderiales | Burkholderiaceae_A_595421 | | |

| | | | | | | | | | | | | | |
|---|---|---|---|---|---|---|---|---|---|---|---|---|---|
| Control UV vs. Control noUV | Control noUV | 20.003 | -41.622 | <0.0001 | 0.0003 | ASV172 | Bacteria | Bacillota_I | Bacilli_A | Lactobacillales | Streptococcaceae | Lactococcus_A_346120 | |
| Control UV vs. Control noUV | Control UV | 13.277 | 37.361 | <0.0001 | 0.0003 | ASV134 | Bacteria | Actinomycetota | Actinomycetes | Mycobacteriales | Micromonosporaceae | | |
| Control UV vs. Control noUV | Control noUV | 81.773 | -24.842 | <0.0001 | 0.0008 | ASV37 | Bacteria | Pseudomonadota | Alphaproteobacteria | Rhizobiales_505101 | Beijerinckiaceae | Methylobacterium | Methylobacterium sp001424705 |
| Control UV vs. Control noUV | Control noUV | 301.191 | -26.722 | <0.0001 | 0.0008 | ASV100 | Bacteria | Actinomycetota | Actinomycetes | Propionibacteriales | Propionibacteriaceae | Cutibacterium | Cutibacterium acnes |
| Control UV vs. Control noUV | Control UV | 78.456 | 23.19 | <0.0001 | 0.001 | ASV61 | Bacteria | Desulfobacterota_G_459546 | Desulfuromonadia | Geobacterales | Geobacteraceae_439684 | | |
| Control UV vs. Control noUV | Control UV | 48.705 | 26.296 | <0.0001 | 0.001 | ASV62 | Bacteria | Pseudomonadota | Alphaproteobacteria | Rhizobiales_505101 | Beijerinckiaceae | Lichenihabitans_504419 | Lichenihabitans ramalinae |
| Control UV vs. Control noUV | Control UV | 37.44 | 22.522 | <0.0001 | 0.0016 | ASV40 | Bacteria | Actinomycetota | Actinomycetes | Actinomycetales | Microbacteriaceae | Amnibacterium_382409 | |
| Control UV vs. Control noUV | Control UV | 52.77 | 23.027 | <0.0001 | 0.0018 | ASV55 | Bacteria | Bacteroidota | Bacteroidia | Cytophagales_B | Hymenobacteraceae | Hymenobacter_910554 | Hymenobacter setariae |
| Control UV vs. Control noUV | Control noUV | 26.974 | -21.894 | 0.0001 | 0.0019 | ASV58 | Bacteria | Pseudomonadota | Alphaproteobacteria | Acetobacterales | Acetobacteraceae | Lichenicoccus | Lichenicoccus roseus |
| Control UV vs. Control noUV | Control UV | 259.261 | 20.733 | 0.0002 | 0.0035 | ASV57 | Bacteria | Pseudomonadota | Gammaproteobacteria | Enterobacterales_737866 | Enterobacteriaceae_A_732334 | Arsenophonus | |
| Control UV vs. Control noUV | Control noUV | 2.695 | -33.248 | 0.0002 | 0.0035 | ASV379 | Bacteria | Acidobacteriota | Terriglobia | Bryobacterales | Bryobacteraceae | KBS-96 | KBS-96 sp000381625 |
| Control UV vs. Control noUV | Control noUV | 91.323 | -32.698 | 0.0003 | 0.004 | ASV351 | Bacteria | Bacillota_I | Bacilli_A | Bacillales_B_306087 | DSM-18226_301387 | | |
| Control UV vs. Control noUV | Control UV | 8.768 | 32.299 | 0.0003 | 0.0044 | ASV75 | Bacteria | Bacteroidota | Bacteroidia | Cytophagales_B | Hymenobacteraceae | Hymenobacter_910554 | Hymenobacter setariae |
| Control UV vs. Control noUV | Control UV | 20.423 | 32.159 | 0.0004 | 0.0044 | ASV252 | Bacteria | Pseudomonadota | Alphaproteobacteria | Rhodobacterales | Rhodobacteraceae | Paracoccus | |
| Control UV vs. Control noUV | Control noUV | 110.455 | -31.351 | 0.0005 | 0.0058 | ASV288 | Bacteria | Actinomycetota | Actinomycetes | Actinomycetales | Micrococcaceae | | |
| Control UV vs. Control noUV | Control noUV | 35.122 | -28.771 | 0.0008 | 0.0079 | ASV47 | Bacteria | Pseudomonadota | Alphaproteobacteria | Rhizobiales_505101 | Beijerinckiaceae | Methylobacterium | Methylobacterium sp001424705 |
| Control UV vs. Control noUV | Control noUV | 263.592 | -30.419 | 0.0007 | 0.0079 | ASV374 | Bacteria | Bacillota_I | Bacilli_A | Bacillales_B_310392 | Bacillaceae_G | Bacillus_A | |
| Control UV vs. Control noUV | Control UV | 24.266 | 21.55 | 0.001 | 0.0099 | ASV81 | Bacteria | Pseudomonadota | Alphaproteobacteria | Acetobacterales | Acetobacteraceae | Lichenicoccus | Lichenicoccus roseus |
| Control UV vs. Control noUV | Control noUV | 23.836 | -29.509 | 0.0011 | 0.0099 | ASV634 | Bacteria | Bacillota_A_368345 | Clostridia_258483 | Oscillospirales | Ethanoligenenaceae | Ruminiclostridium_D | Ruminiclostridium_D cellulosi |
| Control UV vs. Control noUV | Control UV | 2.965 | 29.327 | 0.0011 | 0.0103 | ASV341 | Bacteria | Pseudomonadota | Alphaproteobacteria | | | | |
| Control UV vs. Control noUV | Control noUV | 22.018 | -21.269 | 0.0014 | 0.0104 | ASV63 | Bacteria | Pseudomonadota | Alphaproteobacteria | Sphingomonadales | Sphingomonadaceae_486827 | Sphingomonas_N_483429 | |
| Control UV vs. Control noUV | Control noUV | 4.481 | -28.957 | 0.0013 | 0.0104 | ASV93 | Bacteria | Pseudomonadota | Alphaproteobacteria | Rhizobiales_505101 | Rhizobiaceae | | |
| Control UV vs. Control noUV | Control noUV | 66.071 | -28.94 | 0.0013 | 0.0104 | ASV150 | Bacteria | Actinomycetota | Actinomycetes | Propionibacteriales | Nocardioidaceae | Nocardioides_A_392796 | |

| | | | | | | | | | | | | | |
|---|---|---|---|---|---|---|---|---|---|---|---|---|---|
| Control UV vs. Control noUV | Control noUV | 18.825 | -28.911 | 0.0013 | 0.0104 | ASV850 | Bacteria | Bacteroidota | Bacteroidia | Bacteroidales | Bacteroidaceae | Prevotella | |
| Control UV vs. Control noUV | Control UV | 13.048 | 21.341 | 0.0016 | 0.0118 | ASV107 | Bacteria | Pseudomonadota | Alphaproteobacteria | Acetobacterales | Acetobacteraceae | | |
| Control UV vs. Control noUV | Control noUV | 51.64 | -17.965 | 0.0021 | 0.0149 | ASV101 | Bacteria | Pseudomonadota | Gammaproteobacteria | Burkholderiales | Burkholderiaceae_A_595421 | | |
| Control UV vs. Control noUV | Control UV | 30.609 | 27.588 | 0.0022 | 0.0151 | ASV115 | Bacteria | Actinomycetota | Actinomycetes | Actinomycetales | Micrococcaceae | Micrococcus | Micrococcus luteus |
| Control UV vs. Control noUV | Control UV | 7.758 | 27.093 | 0.0026 | 0.0174 | ASV69 | Bacteria | Pseudomonadota | Alphaproteobacteria | Sphingomonadales | Sphingomonadaceae_486827 | Sphingomonas_L_486704 | Sphingomonas_L_486704 olei |
| Control UV vs. Control noUV | Control noUV | 14.056 | -18.989 | 0.0029 | 0.0187 | ASV91 | Bacteria | Pseudomonadota | Alphaproteobacteria | Rhizobiales_505101 | | | |
| Control UV vs. Control noUV | Control UV | 14.311 | 22.416 | 0.0031 | 0.0192 | ASV105 | Bacteria | Actinomycetota | Actinomycetes | Actinomycetales | Microbacteriaceae | | |
| Control UV vs. Control noUV | Control noUV | 46.867 | -20.421 | 0.0034 | 0.0203 | ASV65 | Bacteria | Bacteroidota | Bacteroidia | Sphingobacteriales | Sphingobacteriaceae | Pedobacter_B_887417 | |
| Control UV vs. Control noUV | Control noUV | 3.112 | -25.525 | 0.0046 | 0.027 | ASV206 | Bacteria | Actinomycetota | Actinomycetes | Actinomycetales | Dermatophilaceae | | |
| Control UV vs. Control noUV | Control noUV | 14.928 | -18.321 | 0.0051 | 0.0288 | ASV117 | Bacteria | Pseudomonadota | Alphaproteobacteria | Caulobacterales | Caulobacteraceae | | |
| Control UV vs. Control noUV | Control UV | 23.045 | 20.975 | 0.0055 | 0.0293 | ASV76 | Bacteria | Bacteroidota | Bacteroidia | Sphingobacteriales | Sphingobacteriaceae | Mucilaginibacter_A | |
| Control UV vs. Control noUV | Control UV | 9.238 | 21.789 | 0.0054 | 0.0293 | ASV86 | Bacteria | Pseudomonadota | Alphaproteobacteria | Caulobacterales | Caulobacteraceae | Caulobacter_487784 | |
| Control UV vs. Control noUV | Control UV | 12.357 | 22.848 | 0.0081 | 0.042 | ASV125 | Bacteria | Actinomycetota | Actinomycetes | Streptomycetales_400645 | Streptomycetaceae_400641 | Streptomyces_400150 | |

**Supplementary Table 3b**. Differentially abundant fungal ASVs across treatments (shown in Fig. 2, 3 of the main text). Statistical parameters were calculated using *DESeq2* .

| Contrast | Direction | Base Mean | Log2 Fold Change | *P* | *Padj* | ASV | Kingdom | Phylum | Class | Order | Family | Genus | Species |
|---|---|---|---|---|---|---|---|---|---|---|---|---|---|
| Sun start vs. Shade start | Shade start | 2.774 | -23.27 | 0.0005 | 0.0097 | f_ASV99 | Fungi | Chytridiomycota | Spizellomycetes | Spizellomycetales | Powellomycetaceae | Thoreauomyces | Thoreauomyces_sp |
| Sun start vs. Shade start | Shade start | 3.697 | -23.4 | 0.0004 | 0.0097 | f_ASV209 | Fungi | Fungi_phy_Incertae_sedis | Fungi_cls_Incertae_sedis | Fungi_ord_Incertae_sedis | Fungi_fam_Incertae_sedis | Fungi_gen_Incertae_sedis | Fungi_sp |
| Sun start vs. Shade start | Shade start | 5.239 | -24.159 | 0.0003 | 0.0097 | f_ASV216 | Fungi | Fungi_phy_Incertae_sedis | Fungi_cls_Incertae_sedis | Fungi_ord_Incertae_sedis | Fungi_fam_Incertae_sedis | Fungi_gen_Incertae_sedis | Fungi_sp |
| Sun start vs. Shade start | Shade start | 5.65 | -23.968 | 0.0003 | 0.0097 | f_ASV480 | Fungi | Ascomycota | Dothideomycetes | Mycosphaerellales | Teratosphaeriaceae | Microcyclospora | Microcyclospora_quercina |
| Sun start vs. Shade start | Shade start | 6.416 | -24.181 | 0.0003 | 0.0097 | f_ASV1119 | Fungi | Basidiomycota | Agaricostilbomycetes | Agaricostilbomycetes_ord_Incertae_sedis | Agaricostilbomycetes_fam_Incertae_sedis | Agaricostilbomycetes_gen_Incertae_sedis | Agaricostilbomycetes_sp |
| Sun start vs. Shade start | Shade start | 30.758 | -24.878 | 0.0002 | 0.0097 | f_ASV1254 | Fungi | Ascomycota | Dothideomycetes | Dothideales | Dothideaceae | Neodothiora | Neodothiora_populina |
| Sun start vs. Shade start | Shade start | 12.806 | -25.39 | 0.0001 | 0.0097 | f_ASV1543 | Fungi | Ascomycota | Taphrinomycetes | Taphrinales | Taphrinaceae | Taphrina | Taphrina_carpini |
| Sun start vs. Shade start | Shade start | 5.285 | -24.169 | 0.0003 | 0.0097 | f_ASV1792 | Fungi | Ascomycota | Ascomycota_cls_Incertae_sedis | Ascomycota_ord_Incertae_sedis | Ascomycota_fam_Incertae_sedis | Ascomycota_gen_Incertae_sedis | Ascomycota_sp |
| Sun start vs. Shade start | Shade start | 33.874 | -23.217 | 0.0005 | 0.0097 | f_ASV2326 | Fungi | Basidiomycota | Microbotryomycetes | Sporidiobolales | Sporidiobolaceae | Sporobolomyces | Sporobolomyces_roseus |
| Sun start vs. Shade start | Sun start | 8.377 | 23.476 | 0.0004 | 0.0097 | f_ASV452 | Fungi | Ascomycota | Dothideomycetes | Pleosporales | Pleosporaceae | Alternaria | Alternaria_metachromatica |
| Sun start vs. Shade start | Sun start | 19.002 | 23.288 | 0.0004 | 0.0097 | f_ASV1269 | Fungi | Ascomycota | Dothideomycetes | Dothideales | Saccotheciaceae | Aureobasidium | Aureobasidium_pullulans |
| Sun start vs. Shade start | Sun start | 7.504 | 23.695 | 0.0004 | 0.0097 | f_ASV1534 | Fungi | Ascomycota | Taphrinomycetes | Taphrinales | Taphrinaceae | Taphrina | Taphrina_sadebeckii |
| Sun start vs. Shade start | Sun start | 52.756 | 26.14 | <0.0001 | 0.0097 | f_ASV2606 | Fungi | Basidiomycota | Tremellomycetes | Filobasidiales | Filobasidiaceae | Filobasidium | Filobasidium_chernovii |
| Sun start vs. Shade start | Shade start | 2.725 | -22.921 | 0.0006 | 0.0103 | f_ASV2765 | Fungi | Ascomycota | Eurotiomycetes | Chaetothyriales | Herpotrichiellaceae | Cladophialophora | Cladophialophora_sp |
| Sun start vs. Shade start | Shade start | 225.28 | -22.85 | 0.0006 | 0.0103 | f_ASV4323 | Fungi | Ascomycota | Sordariomycetes | Hypocreales | Nectriaceae | Fusarium | Fusarium_oxysporum |
| Sun start vs. Shade start | Sun start | 2.92 | 21.913 | 0.001 | 0.0164 | f_ASV1880 | Fungi | Ascomycota | Saccharomycetes | Saccharomycetales | Debaryomycetaceae | Kurtzmaniella | Kurtzmaniella_sp |
| Sun start vs. Shade start | Sun start | 2.573 | 21.274 | 0.0013 | 0.0214 | f_ASV1885 | Fungi | Ascomycota | Saccharomycetes | Saccharomycetales | Debaryomycetaceae | Debaryomyces | Debaryomyces_udenii |
| Sun UV vs. Sun noUV | Sun UV | 266.18 | 30.935 | <0.0001 | <0.0001 | f_ASV459 | Fungi | Ascomycota | Dothideomycetes | Pleosporales | Pleosporaceae | Alternaria | Alternaria_metachromatica |
| Sun UV vs. Sun noUV | Sun UV | 38.564 | 29.273 | <0.0001 | <0.0001 | f_ASV454 | Fungi | Ascomycota | Dothideomycetes | Pleosporales | Pleosporaceae | Alternaria | Alternaria_metachromatica |
| Sun UV vs. Sun noUV | Sun UV | 2.92 | 31.47 | <0.0001 | <0.0001 | f_ASV1880 | Fungi | Ascomycota | Saccharomycetes | Saccharomycetales | Debaryomycetaceae | Kurtzmaniella | Kurtzmaniella_sp |
| Sun UV vs. Sun noUV | Sun noUV | 21.087 | -25.81 | <0.0001 | <0.0001 | f_ASV1455 | Fungi | Ascomycota | Taphrinomycetes | Taphrinales | Taphrinaceae | Taphrina | Taphrina_sp |
| Sun UV vs. Sun noUV | Sun UV | 38.221 | 29.2 | <0.0001 | <0.0001 | f_ASV2327 | Fungi | Basidiomycota | Microbotryomycetes | Sporidiobolales | Sporidiobolaceae | Sporobolomyces | Sporobolomyces_roseus |

| | | | | | | | | | | | | | |
|---|---|---|---|---|---|---|---|---|---|---|---|---|---|
| Sun UV vs. Sun noUV | Sun UV | 0.946 | 26.628 | <0.0001 | <0.0001 | f_ASV2040 | Fungi | Fungi_phy_Incertae_sedis | Fungi_cls_Incertae_sedis | Fungi_ord_Incertae_sedis | Fungi_fam_Incertae_sedis | Fungi_gen_Incertae_sedis | Fungi_sp |
| Shade UV vs. Shade noUV | Shade noUV | 49.184 | -26.334 | <0.0001 | <0.0001 | f_ASV419 | Fungi | Ascomycota | Dothideomycetes | Pleosporales | Pleosporaceae | Alternaria | Alternaria_sp |
| Shade UV vs. Shade noUV | Shade noUV | 149.17 | -25.754 | <0.0001 | <0.0001 | f_ASV201 | Fungi | Fungi_phy_Incertae_sedis | Fungi_cls_Incertae_sedis | Fungi_ord_Incertae_sedis | Fungi_fam_Incertae_sedis | Fungi_gen_Incertae_sedis | Fungi_sp |
| Shade UV vs. Shade noUV | Shade UV | 60.507 | 36.275 | <0.0001 | <0.0001 | f_ASV4325 | Fungi | Ascomycota | Sordariomycetes | Hypocreales | Nectriaceae | Fusarium | Fusarium_oxysporum |
| Shade UV vs. Shade noUV | Shade UV | 60.188 | 35.973 | <0.0001 | <0.0001 | f_ASV974 | Fungi | Fungi_phy_Incertae_sedis | Fungi_cls_Incertae_sedis | Fungi_ord_Incertae_sedis | Fungi_fam_Incertae_sedis | Fungi_gen_Incertae_sedis | Fungi_sp |
| Shade UV vs. Shade noUV | Shade UV | 236.71 | 21.697 | <0.0001 | <0.0001 | f_ASV1258 | Fungi | Ascomycota | Dothideomycetes | Dothideales | Saccotheciaceae | Aureobasidium | Aureobasidium_pullulans |
| Shade UV vs. Shade noUV | Shade noUV | 25.135 | -30.131 | <0.0001 | <0.0001 | f_ASV2025 | Fungi | Ascomycota | Eurotiomycetes | Eurotiales | Aspergillaceae | Aspergillus | Aspergillus_fumigatus |
| Shade UV vs. Shade noUV | Shade noUV | 11.503 | -29.348 | <0.0001 | <0.0001 | f_ASV965 | Fungi | Fungi_phy_Incertae_sedis | Fungi_cls_Incertae_sedis | Fungi_ord_Incertae_sedis | Fungi_fam_Incertae_sedis | Fungi_gen_Incertae_sedis | Fungi_sp |
| Shade UV vs. Shade noUV | Shade UV | 30.758 | 29.095 | <0.0001 | <0.0001 | f_ASV1254 | Fungi | Ascomycota | Dothideomycetes | Dothideales | Dothideaceae | Neodothiora | Neodothiora_populina |
| Shade UV vs. Shade noUV | Shade noUV | 19.002 | -28.032 | <0.0001 | <0.0001 | f_ASV1269 | Fungi | Ascomycota | Dothideomycetes | Dothideales | Saccotheciaceae | Aureobasidium | Aureobasidium_pullulans |
| Shade UV vs. Shade noUV | Shade noUV | 26.682 | -26.692 | <0.0001 | <0.0001 | f_ASV1049 | Fungi | Ascomycota | | | | | |
| Shade UV vs. Shade noUV | Shade UV | 3.476 | 26.312 | <0.0001 | <0.0001 | f_ASV1001 | Fungi | Ascomycota | Sordariomycetes | Hypocreales | Nectriaceae | Nectria | Nectria_ulmicola |
| Shade UV vs. Shade noUV | Shade noUV | 52.579 | -22.935 | <0.0001 | <0.0001 | f_ASV850 | Fungi | | | | | | |
| Shade UV vs. Shade noUV | Shade UV | 2.573 | 23.732 | <0.0001 | <0.0001 | f_ASV1885 | Fungi | Ascomycota | Saccharomycetes | Saccharomycetales | Debaryomycetaceae | Debaryomyces | Debaryomyces_udenii |
| Sun UV vs. Shade noUV | Sun UV | 236.71 | 26.232 | <0.0001 | <0.0001 | f_ASV1258 | Fungi | Ascomycota | Dothideomycetes | Dothideales | Saccotheciaceae | Aureobasidium | Aureobasidium_pullulans |
| Sun UV vs. Shade noUV | Shade noUV | 149.17 | -25.205 | <0.0001 | <0.0001 | f_ASV201 | Fungi | Fungi_phy_Incertae_sedis | Fungi_cls_Incertae_sedis | Fungi_ord_Incertae_sedis | Fungi_fam_Incertae_sedis | Fungi_gen_Incertae_sedis | Fungi_sp |
| Sun UV vs. Shade noUV | Shade noUV | 49.184 | -24.94 | <0.0001 | <0.0001 | f_ASV419 | Fungi | Ascomycota | Dothideomycetes | Pleosporales | Pleosporaceae | Alternaria | Alternaria_sp |
| Sun UV vs. Shade noUV | Sun UV | 33.874 | 33.935 | <0.0001 | <0.0001 | f_ASV2326 | Fungi | Basidiomycota | Microbotryomycetes | Sporidiobolales | Sporidiobolaceae | Sporobolomyces | Sporobolomyces_roseus |
| Sun UV vs. Shade noUV | Shade noUV | 25.135 | -29.517 | <0.0001 | <0.0001 | f_ASV2025 | Fungi | Ascomycota | Eurotiomycetes | Eurotiales | Aspergillaceae | Aspergillus | Aspergillus_fumigatus |
| Sun UV vs. Shade noUV | Shade noUV | 11.503 | -29.192 | <0.0001 | <0.0001 | f_ASV965 | Fungi | Fungi_phy_Incertae_sedis | Fungi_cls_Incertae_sedis | Fungi_ord_Incertae_sedis | Fungi_fam_Incertae_sedis | Fungi_gen_Incertae_sedis | Fungi_sp |
| Sun UV vs. Shade noUV | Sun UV | 38.564 | 24.585 | <0.0001 | <0.0001 | f_ASV454 | Fungi | Ascomycota | Dothideomycetes | Pleosporales | Pleosporaceae | Alternaria | Alternaria_metachromatica |
| Sun UV vs. Shade noUV | Shade noUV | 19.002 | -27.276 | <0.0001 | <0.0001 | f_ASV1269 | Fungi | Ascomycota | Dothideomycetes | Dothideales | Saccotheciaceae | Aureobasidium | Aureobasidium_pullulans |
| Sun UV vs. Shade noUV | Sun UV | 38.221 | 25.587 | <0.0001 | <0.0001 | f_ASV2327 | Fungi | Basidiomycota | Microbotryomycetes | Sporidiobolales | Sporidiobolaceae | Sporobolomyces | Sporobolomyces_roseus |
| Sun UV vs. Shade noUV | Shade noUV | 26.682 | -25.444 | <0.0001 | <0.0001 | f_ASV1049 | Fungi | Ascomycota | | | | | |
| Sun UV vs. Shade noUV | Shade noUV | 52.579 | -24.239 | <0.0001 | <0.0001 | f_ASV850 | Fungi | | | | | | |
| Sun UV vs. Shade noUV | Sun UV | 0.946 | 25.092 | <0.0001 | <0.0001 | f_ASV2040 | Fungi | Fungi_phy_Incertae_sedis | Fungi_cls_Incertae_sedis | Fungi_ord_Incertae_sedis | Fungi_fam_Incertae_sedis | Fungi_gen_Incertae_sedis | Fungi_sp |
| Control UV vs. Control noUV | Control UV | 3880.5 | 18.112 | <0.0001 | 0.0006 | f_ASV338 | Fungi | | | | | | |

**Supplementary Table 4.** Results of ANOVA (**a**) and post hoc Tukey's HSD (**b**) testing the effect of inoculation type (origin of microbial inoculum: sun /shade) and UV exposure (UV+ / UV) on plant performance (Fig. 4 in the main text). Significant results are indicated in bold.

a) Two-factorial ANOVA

| **Plant trait** | **Inoculation type** | | **UV exposure** | | **Inoculation type x UV exposure** | |
|---|---|---|---|---|---|---|
| | $F_{1,52}$ | *P* | $F_{1,52}$ | *P* | $F_{6,138}$ | *P* |
| NDVI | 2.500 | 0.120 | 4.566 | **0.037** | 5.691 | **0.021** |
| Δ $H_{max}$ | 1.116 | 0.296 | 5.378 | **0.024** | 6.230 | **0.016** |

b) Tukey's HSD test

| **Plant trait** | **Comparison** | **Difference [95% CI]** | **Adjusted *P*** |
|---|---|---|---|
| NDVI | Sun_UV- vs. Shade_UV- | -0.035 [-0.094, 0.024] | 0.398 |
| NDVI | Shade_UV+ vs. Shade_UV- | -0.048 [-0.107, 0.012] | 0.155 |
| NDVI | Sun_UV+ vs. Shade_UV- | -0.008 [-0.067, 0.051] | 0.986 |
| NDVI | Shade_UV+ vs. Sun_UV- | -0.012 [-0.071, 0.047] | 0.945 |
| NDVI | Sun_UV+ vs. Sun_UV- | 0.028 [-0.032, 0.087] | 0.607 |
| NDVI | Sun_UV+ vs. Shade_UV+ | 0.040 [-0.019, 0.099] | 0.288 |
| Δ $H_{max}$ | Sun_UV- vs. Shade_UV- | -8.498 [-29.845, 12.849] | 0.717 |
| Δ $H_{max}$ | Shade_UV+ vs. Shade_UV- | -18.653 [-40.000, 2.694] | 0.107 |
| Δ $H_{max}$ | Sun_UV+ vs. Shade_UV- | 1.238 [-20.109, 22.585] | 0.999 |
| Δ $H_{max}$ | Shade_UV+ vs. Sun_UV- | -10.155 [-31.502, 11.192] | 0.590 |
| Δ $H_{max}$ | Sun_UV+ vs. Sun_UV- | 9.736 [-11.611, 31.083] | 0.623 |
| Δ $H_{max}$ | Sun_UV+ vs. Shade_UV+ | 19.891 [-1.456, 41.238] | 0.076 |

**Supplementary Table 5.** Bacterial and fungal ASVs significantly correlated with plant traits within the correlation-based networks (Fig. 5 in the main text).

| **ASV ID** | **Plant trait** | ***rho*** | ***P*** | **Kingdom** | **Phylum** | **Class** | **Order** | **Family** | **Genus** | **Species** |
|---|---|---|---|---|---|---|---|---|---|---|
| ASV12 | Height_max_difference | 0.411 | 0.0371 | Bacteria | Pseudomonadota | Alphaproteobacteria | Sphingomonadales | Sphingomonadaceae_486827 | Sphingomonas_L_486704 | |
| ASV12 | Height_max_difference | 0.425 | 0.034 | Bacteria | Pseudomonadota | Alphaproteobacteria | Sphingomonadales | Sphingomonadaceae_486827 | Sphingomonas_L_486704 | |
| ASV14 | Light_penetration_depth | -0.448 | 0.0247 | Bacteria | Actinomycetota | Actinomycetes | Propionibacteriales | Propionibacteriaceae | Friedmanniella | |
| ASV168 | greenness_average | 0.401 | 0.0472 | Bacteria | Pseudomonadota | Alphaproteobacteria | Rhizobiales_505101 | Beijerinckiaceae | Methylobacterium | Methylobacterium radiotolerans |
| ASV17 | greenness_average | -0.418 | 0.0375 | Bacteria | Actinomycetota | Actinomycetes | Actinomycetales | Microbacteriaceae | Curtobacterium | |
| ASV17 | Digital_biomass_difference | -0.413 | 0.0404 | Bacteria | Actinomycetota | Actinomycetes | Actinomycetales | Microbacteriaceae | Curtobacterium | |
| ASV19 | hue_average | -0.471 | 0.0152 | Bacteria | Actinomycetota | Actinomycetes | Propionibacteriales | Propionibacteriaceae | Cutibacterium | Cutibacterium acnes |
| ASV19 | PSRI_average | -0.426 | 0.0336 | Bacteria | Actinomycetota | Actinomycetes | Propionibacteriales | Propionibacteriaceae | Cutibacterium | Cutibacterium acnes |
| ASV19 | NPCI_average | 0.531 | 0.0052 | Bacteria | Actinomycetota | Actinomycetes | Propionibacteriales | Propionibacteriaceae | Cutibacterium | Cutibacterium acnes |
| ASV19 | PSRI_average | 0.434 | 0.0268 | Bacteria | Actinomycetota | Actinomycetes | Propionibacteriales | Propionibacteriaceae | Cutibacterium | Cutibacterium acnes |
| ASV25 | Light_penetration_depth | -0.454 | 0.0227 | Bacteria | Pseudomonadota | Alphaproteobacteria | Acetobacterales | Acetobacteraceae | Pseudoroseomonas | |
| ASV26 | Height_max_difference | -0.398 | 0.049 | Bacteria | Pseudomonadota | Gammaproteobacteria | Pseudomonadales_A_650611 | Pseudomonadaceae | | |
| ASV273 | Light_penetration_depth | -0.41 | 0.0419 | Bacteria | Pseudomonadota | Alphaproteobacteria | Rhizobiales_505101 | Xanthobacteraceae | | |
| ASV273 | Digital_biomass_difference | -0.474 | 0.0166 | Bacteria | Pseudomonadota | Alphaproteobacteria | Rhizobiales_505101 | Xanthobacteraceae | | |
| ASV273 | Leaf_area_index | -0.457 | 0.0215 | Bacteria | Pseudomonadota | Alphaproteobacteria | Rhizobiales_505101 | Xanthobacteraceae | | |
| ASV273 | Leaf_area | -0.404 | 0.045 | Bacteria | Pseudomonadota | Alphaproteobacteria | Rhizobiales_505101 | Xanthobacteraceae | | |
| ASV28 | hue_average | 0.409 | 0.0378 | Bacteria | Pseudomonadota | Gammaproteobacteria | Enterobacterales_737866 | | | |
| ASV28 | Height_max_difference | 0.548 | 0.0037 | Bacteria | Pseudomonadota | Gammaproteobacteria | Enterobacterales_737866 | | | |
| ASV28 | PSRI_average | -0.436 | 0.026 | Bacteria | Pseudomonadota | Gammaproteobacteria | Enterobacterales_737866 | | | |
| ASV41 | Leaf_area | 0.477 | 0.0137 | Bacteria | Pseudomonadota | Gammaproteobacteria | Pseudomonadales_A_660879 | Moraxellaceae | Moraxella_A | |
| ASV41 | hue_average | 0.448 | 0.0216 | Bacteria | Pseudomonadota | Gammaproteobacteria | Pseudomonadales_A_660879 | Moraxellaceae | Moraxella_A | |
| ASV41 | Light_penetration_depth | 0.567 | 0.0025 | Bacteria | Pseudomonadota | Gammaproteobacteria | Pseudomonadales_A_660879 | Moraxellaceae | Moraxella_A | |
| ASV41 | Digital_biomass_difference | 0.489 | 0.0113 | Bacteria | Pseudomonadota | Gammaproteobacteria | Pseudomonadales_A_660879 | Moraxellaceae | Moraxella_A | |
| ASV41 | NPCI_average | -0.461 | 0.0177 | Bacteria | Pseudomonadota | Gammaproteobacteria | Pseudomonadales_A_660879 | Moraxellaceae | Moraxella_A | |

| | | | | | | | | | | |
|---|---|---|---|---|---|---|---|---|---|---|
| ASV41 | Leaf_area_index | 0.47 | 0.0155 | Bacteria | Pseudomonadota | Gammaproteobacteria | Pseudomonadales_A_660879 | Moraxellaceae | Moraxella_A | |
| ASV41 | Height_max_difference | 0.445 | 0.0229 | Bacteria | Pseudomonadota | Gammaproteobacteria | Pseudomonadales_A_660879 | Moraxellaceae | Moraxella_A | |
| ASV41 | NDVI_average | 0.425 | 0.0306 | Bacteria | Pseudomonadota | Gammaproteobacteria | Pseudomonadales_A_660879 | Moraxellaceae | Moraxella_A | |
| ASV44 | Light_penetration_depth | 0.409 | 0.0426 | Bacteria | Pseudomonadota | Alphaproteobacteria | Rhizobiales_505101 | Xanthobacteraceae | | |
| ASV44 | Leaf_area_index | 0.512 | 0.009 | Bacteria | Pseudomonadota | Alphaproteobacteria | Rhizobiales_505101 | Xanthobacteraceae | | |
| ASV44 | Leaf_area | 0.517 | 0.0081 | Bacteria | Pseudomonadota | Alphaproteobacteria | Rhizobiales_505101 | Xanthobacteraceae | | |
| ASV45 | greenness_average | -0.43 | 0.0284 | Bacteria | Pseudomonadota | Gammaproteobacteria | Xanthomonadales | Xanthomonadaceae | | |
| ASV5 | Height_max_difference | -0.484 | 0.0143 | Bacteria | Bacteroidota | Bacteroidia | Cytophagales_B | Hymenobacteraceae | Hymenobacter_910554 | Hymenobacter sp004684095 |
| ASV5 | Light_penetration_depth | -0.649 | 0.0004 | Bacteria | Bacteroidota | Bacteroidia | Cytophagales_B | Hymenobacteraceae | Hymenobacter_910554 | Hymenobacter sp004684095 |
| ASV9 | Leaf_area_index | -0.502 | 0.0105 | Bacteria | Pseudomonadota | Alphaproteobacteria | Rhizobiales_505101 | Xanthobacteraceae | | |
| ASV9 | NDVI_average | -0.397 | 0.0492 | Bacteria | Pseudomonadota | Alphaproteobacteria | Rhizobiales_505101 | Xanthobacteraceae | | |
| ASV9 | Digital_biomass_difference | -0.48 | 0.0152 | Bacteria | Pseudomonadota | Alphaproteobacteria | Rhizobiales_505101 | Xanthobacteraceae | | |
| ASV9 | Light_penetration_depth | -0.414 | 0.0398 | Bacteria | Pseudomonadota | Alphaproteobacteria | Rhizobiales_505101 | Xanthobacteraceae | | |
| ASV9 | Leaf_area | -0.453 | 0.0231 | Bacteria | Pseudomonadota | Alphaproteobacteria | Rhizobiales_505101 | Xanthobacteraceae | | |
| f_ASV1274 | Light_penetration_depth | 0.612 | 0.0009 | Fungi | Ascomycota | Dothideomycetes | Dothideales | Saccotheciaceae | Aureobasidium | Aureobasidium_pullulans |
| f_ASV1274 | Digital_biomass_difference | 0.539 | 0.0045 | Fungi | Ascomycota | Dothideomycetes | Dothideales | Saccotheciaceae | Aureobasidium | Aureobasidium_pullulans |
| f_ASV1274 | Leaf_area | 0.543 | 0.0042 | Fungi | Ascomycota | Dothideomycetes | Dothideales | Saccotheciaceae | Aureobasidium | Aureobasidium_pullulans |
| f_ASV1274 | Height_max_difference | 0.455 | 0.0194 | Fungi | Ascomycota | Dothideomycetes | Dothideales | Saccotheciaceae | Aureobasidium | Aureobasidium_pullulans |
| f_ASV1274 | Leaf_area_index | 0.509 | 0.0079 | Fungi | Ascomycota | Dothideomycetes | Dothideales | Saccotheciaceae | Aureobasidium | Aureobasidium_pullulans |
| f_ASV266 | Height_max_difference | 0.582 | 0.0023 | Fungi | | | | | | |
